# Transcriptional adaptation of sensory neurons to GPCR identity and activity

**DOI:** 10.1101/2021.11.29.468971

**Authors:** Luis Flores Horgue, Alexis Assens, Leon Fodoulian, Leonardo Marconi Archinto, Joël Tuberosa, Alexander Haider, Madlaina Boillat, Alan Carleton, Ivan Rodriguez

**Author notes:** equal contribution (Luis Flores Horgue, Alexis Assens).

## Abstract

Sensory adaptation is critical to extract information from a changing world. Taking advantage of the extensive parallel coding lines present in the olfactory system, we explored the potential variations of neuronal identities before and after olfactory experience. We found that at rest, the transcriptomic profiles of olfactory sensory neuron populations are already highly divergent, specific to the olfactory receptor they express, and are surprisingly associated with the sequence of these latter. These divergent profiles further evolve in response to the environment, as odorant exposure leads to massive reprogramming via the modulation of transcription. Adenylyl cyclase 3, but not other main elements of the olfactory transduction cascade, plays a critical role in this activity-induced transcriptional adaptation. These findings highlight a broad range of sensory neuron identities that are present at rest and that adapt to the experience of the individual, thus providing a novel layer of complexity to sensory coding.

## Introduction

Mammals use various sensory tools to build a representation of the outside world. A precise and robust representation being critical for survival and reproduction, we evolved a considerable number of different receptors able to respond to external stimuli. This is particularly true for chemical recognition, for which most of us benefit from large odorant chemoreceptor gene repertoires, that range in the hundreds in humans and dogs, up to over a thousand in mice and elephants (Bear et al., 2016; Jiang and Matsunami, 2015; Niimura and Nei, 2006). Our understanding of olfactory information coding is based on the expression of a single chemoreceptor gene, stochastically chosen from a single allele, in each olfactory sensory neuron; this is referred to as singular expression (Chess et al., 1994; Dalton and Lomvardas, 2015; Mombaerts, 2004; Saraiva et al., 2015). Given the millions of olfactory sensory neurons present in a nasal cavity, large neuronal populations with identical agonist receptivity coexist with other populations that exhibit different response profiles. Olfactory receptors being able to bind various molecules and a volatile to be recognized by different receptors, to any given olfactory stimulus corresponds a specific pattern of activation. The term combinatorial coding has been coined to describe this encoding of chemical identities (Buck, 2004; Malnic et al., 1999). The odorant-dependent activation map is not just a concept, but is directly observable in the olfactory bulb, where axonal projections of olfactory sensory neurons coalesce into neuropil-rich structures, the glomeruli, that are each innervated by a specific sensory population. Following this first relay of olfactory processing and after being transmitted to cortical areas, olfactory information often translates into a percept, a smell. A series of parallel and invariable coding lines, defined by the expressed odorant receptor (and a few guidance molecules that may be differentially expressed between populations), is thus at the heart of the way we understand olfaction.

The ability for neurons to adapt, to respond dynamically to the environment, both during development and later, is critical (West and Greenberg, 2011). Following perturbations of their surroundings, rapid and long-term changes thus occur in cells, which in sensory systems for example lead to sensory adaptation and to compensatory plasticity (Davis, 2006). Various mechanisms answering this need have been selected during evolution, among which the repression or activation of specific genes (Benito and Barco, 2015; West and Greenberg, 2011; Yap and Greenberg, 2018). In the olfactory system, this question has been addressed in the past by various groups (Barber and Coppola, 2015; Cadiou et al., 2014; Coppola and Waggener, 2012; Fischl et al., 2014; Ibarra-Soria et al., 2017; Wang et al., 2017; Zhao et al., 2013). Almost without exception, these approaches have involved the silencing of neuronal activity via naris occlusion or activity mutants, thus adding various confounding factors to odorant-induced activity. Due to a lack of technology available at the time, most of these approaches explored the system at the level of the whole olfactory mucosa, precluding an evaluation of the response to a specific ligand of specific neurons or specific neuronal populations expressing a given odorant receptor. We recently reported that following in vivo odorant exposure of olfactory sensory neurons in the mouse, a decrease in the amount of mRNA encoding for the odorant receptor gene expressed by the activated neurons takes place (von der Weid et al., 2015). Whether this odorant-induced decrease of mRNA concentration is limited to the receptor mRNA, remains to be determined.

Taking advantage of the rare opportunity offered by the mouse olfactory system, composed of large and numerous populations of neurons, each identifiable by the transcription of a specific chemoreceptor gene and thus activable at will by specific ligands, we explored, in the mouse, the organization and the potential modulation of population transcriptional identities. We characterized the transcriptomes of several thousand mouse olfactory sensory neurons and uncovered, at rest, an unexpected variability of profiles, each defined by the differential expression of numerous genes and dependent on the expressed chemoreceptor. Following neuronal activation, we found that a second layer of variability, that relies on adenylyl cyclase 3 and involves transcriptional modulation, is added to this initial diverse landscape.

## Results

### Variable transcriptomes among olfactory sensory neuron populations at rest

The main determinant of an olfactory sensory neuron identity is the olfactory receptor gene it expresses. In addition to this functional characteristic, a few genes have been described to be unequally expressed in different olfactory populations. These latter are however thought to be shared by large numbers of sensory neuron populations (that is populations expressing different odorant receptors), and to be involved in the topographic organization of the olfactory sensory mucosa, in which neurons expressing a given odorant receptor are restricted to specific zones.

As an initial approach to determine potential differences in the identity of the various sensory neurons populating the main olfactory epithelium, we performed a single-cell RNA-seq of dissociated cells extracted from the nasal cavity of 8 week-old male mice (Figure 1A). The data were clustered and visualized on a UMAP plot (Figure 1B). We obtained a total of 15’927 cells, from which mature olfactory sensory neurons (expressing the *Omp* and *Adcy3* genes) were readily distinguishable from immature neurons and non-neuronal cells (Figure 1B and Figure S1A), in agreement with previous results(Fletcher et al., 2017). A total of 9’762 mature olfactory sensory neurons (composed of 811 olfactory sensory populations of at least three neurons (Figure S1D)) expressing the olfactory marker gene *Omp* were then selected, and clustered again. Specific subclusters were observed (Figure 1C and S1B,C), defined by the expression of certain marker genes (*Calb2*, *Cd36* and others), as well as a clear separation between sensory neurons located ventrally (*Nfix*+) and dorsally (*Nqo1+*) in the nasal cavity (olfactory sensory neurons expressing specific odorant receptors are unequally scattered across the nasal epithelium (Buck, 1996)). We then explored the potential transcriptional proximity of neurons expressing the same odorant receptor. Their positions were visualized on the UMAP plot, which revealed a striking grouping of each of the different neuronal populations (Figure 1D,E), ranging from dense clustering of their transcriptomes (such as those expressing *Olfr354* or *Olf553*), to populations exhibiting a larger variance in gene expression (such as *Olfr1183*). To quantify this observation we measured the pairwise transcriptomic distances between pairs of sensory neurons expressing the same receptor and found that neurons expressing the same receptor were significantly more similar transcriptionally than those expressing different receptors (Figure 1E). To determine the potential role played by the expression of the odorant receptors themselves in this clustering, this latter was performed without taking the olfactory receptor expression data into account. Remarkably, the grouping of populations expressing the same chemoreceptor, irrespective of whether the analysis was performed on the whole sensory population or a given subcluster (Figures 1E and S2), was maintained. To explore this olfactory-receptor-associated population specificity, we identified specific genes that were differentially expressed between the different neuronal populations (Figure 1F). These included genes that were either transcribed or whose transcripts were absent in the different populations such as *Cidea*, or that were expressed in a graded manner across most populations, such as *S100a5* (Figure 1F). These transcriptomic profiles were not merely reflecting different general types of sensory neurons, since they were not only observed to be different between subclusters, but also within subclusters (Figure 1G,H). At rest, that is without active olfactory stimulation, a transcriptomic code thus characterizes each odorant receptor-defined population.

**Figure 1.**
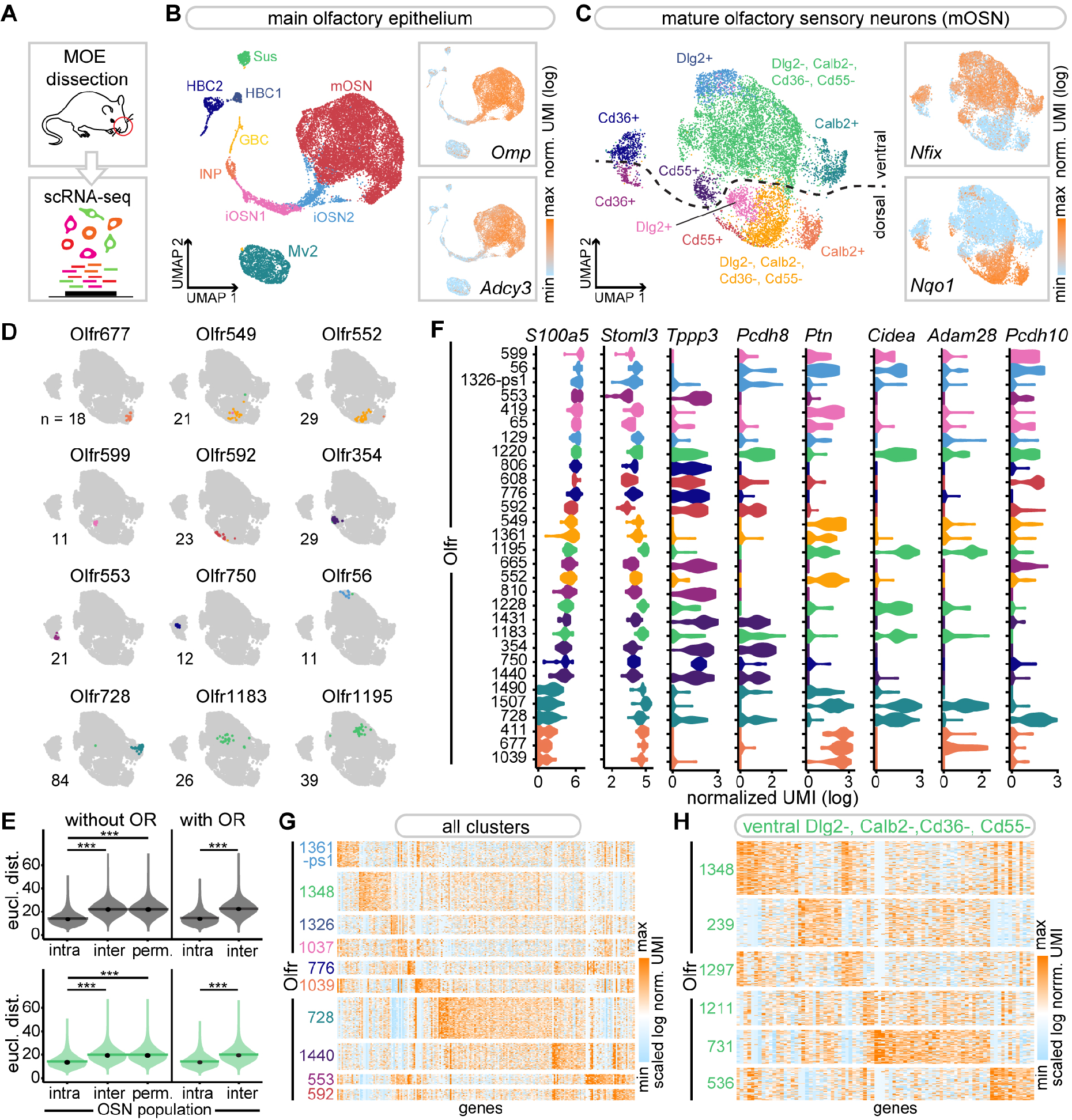
Variable odorant receptor-associated transcriptomes among olfactory sensory populations. (A) Schematic of the approach. (B) Visualization of MOE cell clusters on a *UMAP* plot. *Inset*: normalized expression levels of two mOSN gene markers, *Omp* and *Adcy3*. (C) Visualization of mOSN cell clusters on a *UMAP* plot computed after removing the olfactory receptor gene counts from the data. *Inset*: normalized expression levels of *Nfix* and *Nqo1* (markers of neurons located ventrally and dorsally). (D) Visualization of the dispersion of OSN populations on the UMAP plot shown in (C). The color of each cell indicates the cluster to which the cells pertains. (E) Violin plots showing the density distribution of transcriptomic Euclidean distances (computed on the first 19 PCs) between pairs of OSNs expressing the same receptor (intra), different receptors (inter), and the same receptor after permutation of receptor identities prior to distance calculation (perm.) (n = 1000 permutations, see methods). Data are plotted for all OSNs in (G) or ventral cluster OSNs in (H). Horizontal bars correspond to mean values and dots correspond to median values. \**p*<0.05; \*\**p*<0.01; \*\*\**p*<0.001; Wilcoxon rank sum test. (F) Violin plots showing mOSN population-specific distribution of selected markers (log-normalized UMI). OSN populations are ordered by their mean expression of *S100a5*. The color of each violin plot indicates the cluster to which the majority of the cells from the given population pertain. (G-H) Heatmap representation of the expression levels (scaled log normalized UMI) of the specific gene markers of the largest mOSN populations selected from each cluster (E) or from the ventral Dlg2-, Calb2-, Cd36- and Cd55- clusters (F).

**Figure 2.**
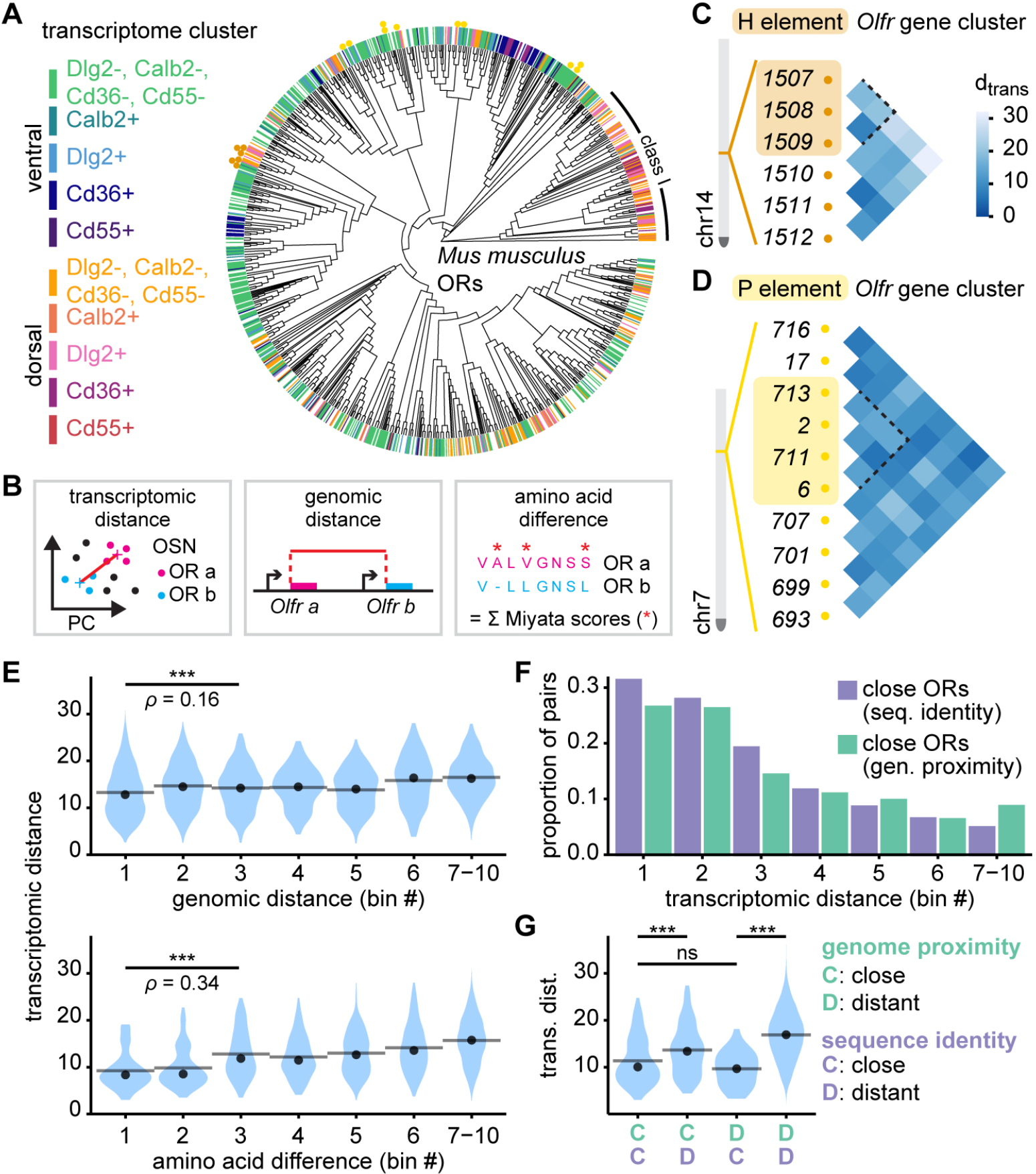
Association between transcriptomic distances and odorant receptor similarity levels. (A) Phylogenetic reconstruction of 1152 mouse odorant receptor proteins. Colored bars at the tree tips show the transcriptomic identity of the olfactory sensory subpopulations expressing a given odorant receptor (labeled receptors are restricted to those detected in three cells or more in our scRNA-seq dataset). Colored dots show odorant receptors belonging to the gene clusters analyzed in C and D, (labeled receptors are restricted to those detected in 10 cells or more in our scRNA-seq dataset). (B) The transcriptomic distances between olfactory sensory populations expressing a given odorant receptor was calculated as the Euclidian distance between centroids in the principal component analysis of mOSN transcriptomes. Genomic distance was calculated as the distance in base pairs between the start codons of Olfr genes; amino acid differences were calculated as the sum of Miyata amino acid replacement scores from an olfactory receptor alignment. Each insertion was scored as the mean replacement score of the additional amino acid. (C) Pairwise transcriptomic distances between olfactory sensory populations characterized by the expression of Olfr genes under the control of the H element on chromosome 14, and their neighbors. H element-regulated Olfr genes are denoted by the box surrounding their names. Cells of the triangular matrix correspond to each pairwise comparison, with rows linking each possible pair. The dotted line encompasses comparisons between olfactory sensory populations expressing each of the H element- regulated Olfr genes. Cells are colored according to the pairwise transcriptomic distance. (D) Same analysis as in (C) but for Olfrs under the control of the P element, on chromosome 7. (E) Distribution of transcriptomic distances per bins of either genomic distances (top) or amino acid differences (bottom), for all pairs of odorant receptor belonging to the same class and the same gene cluster. Spearman’s rank correlation (ρ) and associated p-value between the transcriptomic distance and each of the different metrics was calculated for the pairs included in the three first bins. Transcriptomic distances and genomic distance: ρ=0.16, \*\*\**p*<0.001, n=2761 pairs. Transcriptomic distances and genomic distance: ρ=0.34, \*\*\**p*<0.001, n=233 pairs. (F) Closely related pairs of odorant receptors were identified relative to their genomic distance (95^th^ percentile of the intergenic distances between neighboring Olfr genes) or sequence identity (5^th^ percentile of the pairwise amino acid difference distribution). The proportion of these pairs was determined for each bin of pairwise transcriptomic distances. (G) Transcriptomic distance distribution of four categories of odorant receptor gene pairs defined by their genomic proximity and the sequence identity of their corresponding protein. Close pairs were defined as in F. In terms of genome proximity, distant pairs were defined as belonging to different gene clusters. In terms of sequence identity, distant pairs were defined as higher than the threshold value set for close sequences. \*\*\**p*<10, ns *p*>0.05; Wilcoxon rank test.

### Transcriptomic proximity versus olfactory receptor identity

What may determine the transcriptional distance between two neuronal populations expressing different odorant receptors? Considering olfactory sensory neuron maturation as a series of differentiation steps that at each increment determine more and more specific identities (Fletcher *et al*., 2017; Hanchate et al., 2015), transcriptomic distances between two mature sensory neuron populations may reflect how many of these steps they shared. If transcriptomic identity is largely determined prior to odorant gene choice, the closeness between two populations should be mirrored by their use of the same cis-regulatory element. These elements are known to be necessary for the choice of specific sets of odorant receptor genes and control small clusters of adjacent odorant receptor genes (Bozza et al., 2009; Fuss et al., 2007; Khan et al., 2011; Nishizumi et al., 2007). Alternatively or in addition to this first hypothesis, one could envisage a direct role played by the receptor itself, which may define a basal activity level in neurons for example, and a corresponding transcriptomic profile.

We tested the first hypothesis by evaluating whether olfactory receptor genes sharing a common enhancer are transcriptionally closer to each other than to those under the control of other cis-regulatory elements. We took advantage of two well defined cis-regulatory elements acting on mouse odorant receptor genes (the H and P elements), whose olfactory receptor gene targets are quite dissimilar (Figure 2A,C,D), and have been well described (Bozza *et al*., 2009; Fuss *et al*., 2007; Khan *et al*., 2011; Nishizumi *et al*., 2007). Calculating a centroid-based Euclidian distance between olfactory sensory neuron population transcriptomes (Figure S2B, Figure S4A), we evaluated the similarity between the transcriptomes of sensory neurons subpopulations expressing olfactory receptor genes under the control of the same cis-regulatory elements, as well as their neighbors (Figure 2C,D). No increase in transcriptomic similarity was observed between genes under the control of the H or P elements. To further explore this hypothesis, we took a global approach based on the possible link between the genomic distance separating odorant receptor genes and the difference in transcriptomic identity, the idea being again that since olfactory cis-regulatory elements act on adjacent genes, those located in proximity may also result closer transcriptionally. A weak positive association between genomic distance (Figure 2B) and transcriptomic proximity was observed in the first 30% of the genomic distance value range (Figure 2E, S4B). This association increased when looking at the first 10% (Figure S4C,D). This relationship was more visible when considering different equal width bins of the observed range of transcriptomic distances, in relation to the proportion of pairs of neighbouring genes (Figure 2F).

We then evaluated our second hypothesis, which proposes that the odorant receptor themselves define transcriptomic profiles. We determined the levels of sequence homology between odorant receptors using Miyata scores (Figure 2B, S4E) and tested their potential association with the transcriptomic distances between the corresponding neuronal populations. We found a positive association between transcriptomic proximity and odorant receptor sequence similarity in the first 30% of the amino acid difference value range (Figure 2E, S4E), that was further supported by a clear overrepresentation of similar odorant receptor pairs in sensory populations that are transcriptionally close (Figure 2F, S4E). Given this last observation and knowing that odorant receptor genes tend to duplicate in cis, it is likely that sequence identity is a confounding factor when measuring genomic proximity. To reevaluate whether genomic proximity plays indeed a role in addition to receptor identity, we took advantage of evolutionary accidents that led duplicated odorant receptor genes to land in close vicinity or distantly from their princeps allele. We thus evaluated the potential differences in transcriptional proximity between neuronal populations expressing odorant receptors that are very similar and associated in the genome, and neuronal populations expressing odorant receptors that are very similar but located in different gene clusters. In parallel, we evaluated the transcriptomic distances of neuronal populations expressing odorant receptors that are dissimilar and are associated in the genome (Figure 2G). We found no difference in the average transcriptional distance between neurons expressing similar odorant receptor genes, whether these genes are located in proximity or are distant from each other. On the contrary, we found an increase in transcriptomic distances between dissimilar and similar odorant receptors associated in the genome, supporting our second hypothesis, that is a role played by the odorant receptor identity in transcriptome determination.

### Activity induced transcriptomic modulation in Olrf151- and Olfr16-expressing sensory neurons

Following the identification of specific transcriptomic identities characterizing the different olfactory neuronal populations at rest, we explored their potential evolution after agonist activation. To address this question, we determined the transcriptome of defined and well described neuronal populations following exposure to odorants (Figure 3A). We used two knockin mouse lines, *Olfr151^GFP/GFP^* and *Olfr16^GFP/GFP^*, in which olfactory sensory neurons expressing the *Olfr151* (*M71*) and the *Olfr16* (*MOR23*) odorant receptor genes are modified such that when transcribed, a green fluorophore is coexpressed. These olfactory receptors have very different sequences (Miyata score=273.24), are expressed in different zones of the olfactory epithelium (Figure 3B,D), are expressed in different basoapical layers (Figure 3C-F), and respond to different agonists. 12 live and freely moving mice were exposed for 5 hours to acetophenone and lyral, two known agonists of Olfr151 and Olfr16, respectively (Bozza et al., 2002; Touhara et al., 1999) (Figure 3A). To extract the most possible transcriptomic information, we did not opt here for a scRNA-seq approach but rather for the bulk sequencing of purified neuronal populations. Following exposure, fluorescent *Olfr151*- and *Olfr16*-expressing neurons were isolated by FACS, and their transcriptomes were determined and analyzed. Significant and robust transcriptomic modulations were observed for both olfactory populations after agonist exposure, with 645 and 752 genes upregulated and downregulated respectively for Olfr151, and 419 genes upregulated and 356 downregulated for Olfr16 (Figure 3H-K,L,N,O and Table 1). The foldchange modulation ranged from 0.032 to 1505 for Olfr151 and from 0.056 to 517 for Olfr16 (Figure 3I,K).

**Figure 3.**
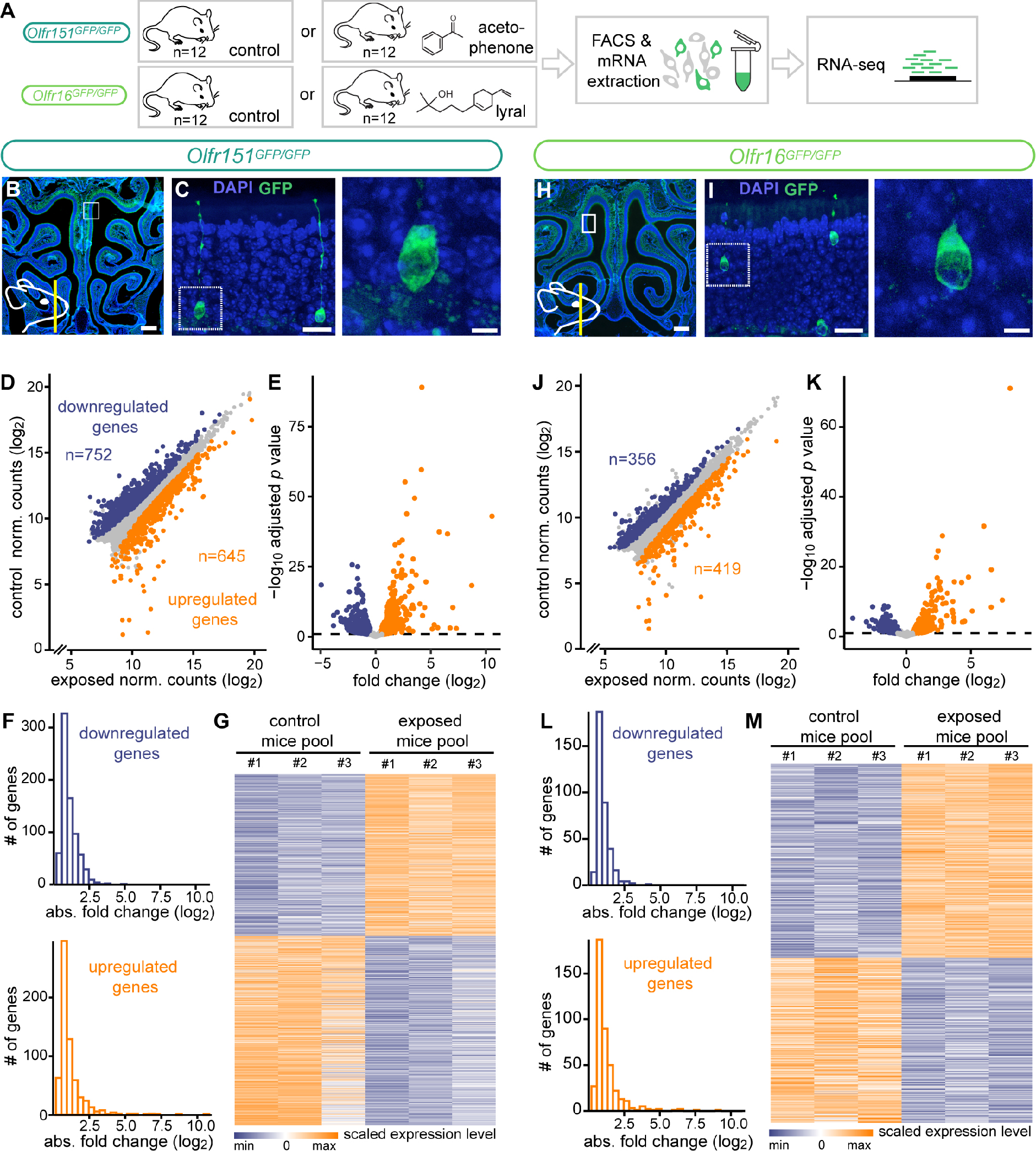
Odorant-induced massive transcriptomic modulations. (A) Schematic of the experiment. After being exposed to their cognate ligand for 5 hours, fluorescent neurons from *Olfr151*^GFP/GFP^ and *Olfr16*^GFP/GFP^ mice were FAC-sorted and total mRNA was sequenced. (B) Coronal section of an *Olfr151*^GFP/GFP^ mouse olfactory epithelium stained with DAPI (blue). The schematic on the lower left indicates the antero-posterior position of the section and the white square the region magnified in (C). Scale bar, 0.5 mm. (C) Section of an *Olfr151*^GFP/GFP^ mouse olfactory epithelium containing GFP-expressing neurons (green, endogenous GFP). The white square highlights the cell magnified in the image on the right. Scale bars, 20 *μ*m (left) and 5 *μ*m (right). (D) Scatter plot showing mean normalized counts resulting from the differential expression analysis (exposed versus non-exposed *Olfr151*^GFP/GFP^ mice). Blue and orange dots: significantly downregulated and upregulated genes, respectively. (E) Volcano plot showing differentially expressed genes (DEGs) between exposed and non-exposed *Olfr151*^GFP/GFP^ mice. The x-axis is the log_2_ scale of the gene expression fold change. Negative values indicate downregulation and positive values upregulation. The y-axis is the minus log10 scale of the adjusted pvalues. Blue and orange dots: significantly downregulated and upregulated genes, respectively. (F) Distribution of the differentially expressed genes between exposed and non exposed *Olfr151*^GFP/GFP^ mice based on their fold change (log_2_). (G) Heatmap showing differentially expressed genes between exposed and non-exposed *Olfr151*^GFP/GFP^ mice. (H) Coronal section of an *Olfr16*^GFP/GFP^ mouse olfactory epithelium stained with DAPI (blue). The schematic on the lower left indicates the antero-posterior position of the section and the white square the region magnified in (C). Scale bar, 0.5 mm. (I) Section of an *Olfr16*^GFP/GFP^ mouse olfactory epithelium containing GFP-expressing neurons (green, endogenous GFP). The white square highlights the cell magnified in the image on the right. Scale bars, 20 *μ*m (left) and 5 *μ*m (right). (J) Scatter plot showing mean normalized counts resulting from the differential expression analysis (exposed versus non-exposed *Olfr16*^GFP/GFP^ mice). Blue and orange dots: significantly downregulated and upregulated genes, respectively. (K) Volcano plot showing differentially expressed genes (DEGs) between exposed and non-exposed *Olfr16*^GFP/GFP^ mice. The x-axis is the log_2_ scale of the gene expression fold change. Negative values indicate downregulation and positive values upregulation. The y-axis is the minus log10 scale of the adjusted pvalues. Blue and orange dots: significantly downregulated and upregulated genes, respectively. (L) Distribution of the differentially expressed genes between exposed and non exposed *Olfr16*^GFP/GFP^ mice based on their fold change (log_2_). (M) Heatmap showing the differentially expressed genes between exposed and non-exposed *Olfr16*^GFP/GFP^ mice.

### Common activity-induced transcriptomic adaptation in Olfr151- and Olfr16-expressing sensory neurons

Taking advantage of our double approach and to potentially extract general rules, we compared the transcriptomic modulations of the Olfr151 and Olfr16 populations after agonist exposure (Figure 4A). In accordance to our initial observations describing significant transcriptomic distances between populations expressing different receptors, we first observed that the transcriptomes of *Olfr151*- and *Olfr16*- expressing neurons were significantly dissimilar (882 genes were differentially expressed between the two neuron populations, Figure 4B-E). We then compared the activity-induced modulated genes between the Olfr151 and Olfr16 populations (Figure 4F-H), that without surprise, showed again a significant difference between populations. A principal component analysis of the Olfr151 and Olfr16 transcriptomic sets, before and after agonist exposure, showed 39% of variance explained by cell identity, and 20% of variance by activity. The overlap between the activity-induced responses of the Olf151 and Olf16 populations was further explored by comparing their potentially common down and upregulated genes. A significant proportion of these genes were shared, 215 (51.3%) and 134 (37.6%) of them being commonly upregulated and downregulated, respectively (Figure 4I). In both Olfr151- and Olfr16- activated populations, the dispersion of downregulated genes decreased, while it increased for upregulated genes; this was also true for the genes shared by both populations (Figure S5). Finally, we aimed to functionally interpret our data by attributing Gene Ontology (GO) terms to the activity-dependent modulated genes common to the Olfr151 and Olfr16 populations. Among the very significantly enriched terms, we found signaling and G-protein-coupled receptor activity in the GO biological processes terms, molecular transducer activity in the GO molecular functions category, and plasma membrane and cell periphery in the GO cellular components category (Figure 4J and Table 2). A pattern thus emerged, pointing to actors of the transduction cascade being modulated by agonist exposure.

**Figure 4.**
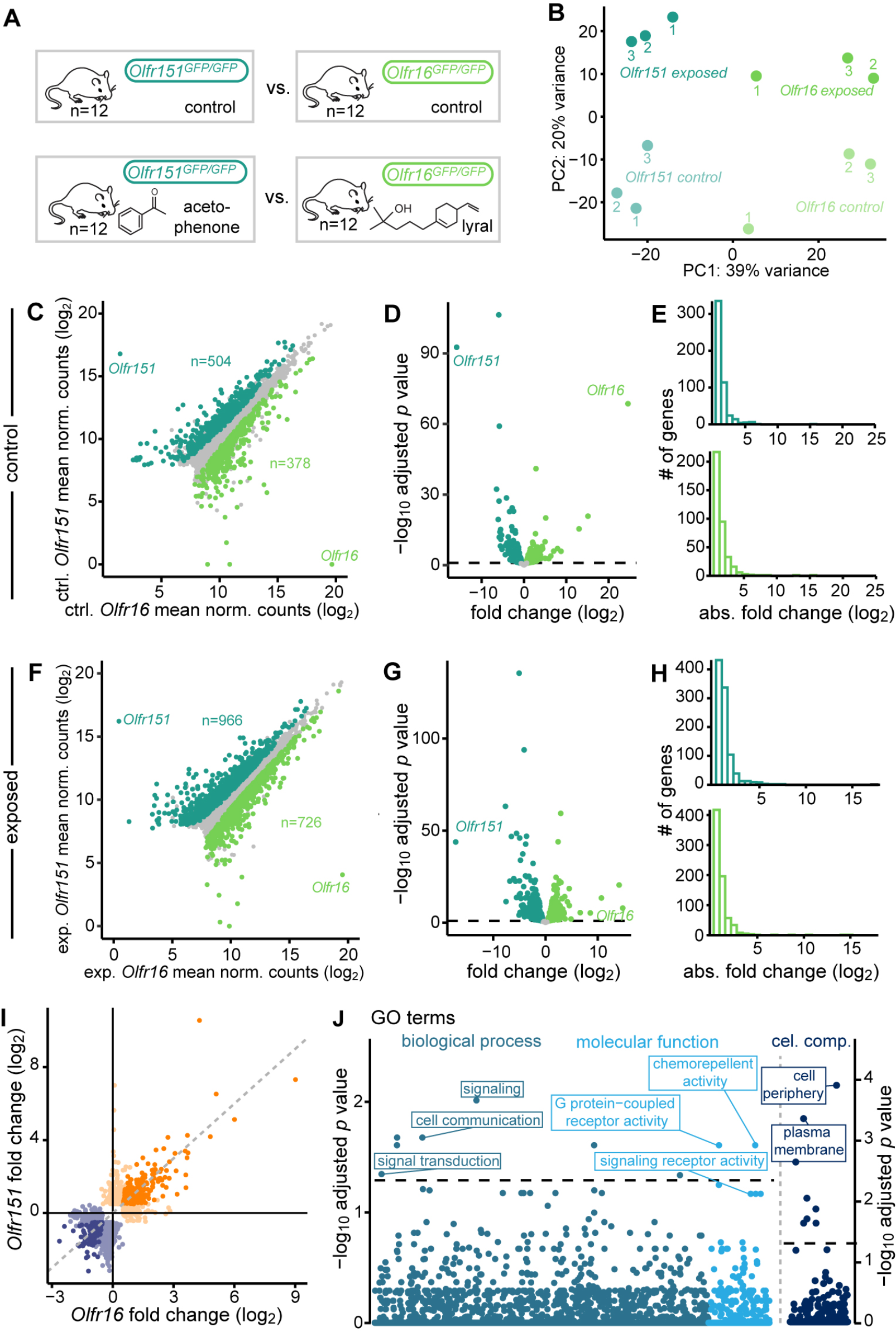
Shared odorant-induced transcriptomic modulations between different olfactory populations. (A) Schematic of the experiment. After being exposed to their cognate ligand, fluorescent neurons from *Olfr151*^GFP/GFP^ and *Olfr16*^GFP/GFP^ mice were FAC-sorted and total mRNA was sequenced. (B) Two dimensional PCA representing the differences in gene expression between two populations of olfactory sensory neurons in their basal state (non-exposed) and after exposure to their cognate ligand. Each dot represents a pool of 4 mice. (C) Scatter plot showing the differential expression analysis between *Olfr15-* and *Olfr16-* expressing neurons in their basal state. Dark and light green dots: genes expressed significantly higher in *Olfr15*- and in *Olfr16*-expressing neurons, respectively. (D) Volcano plot showing differentially expressed genes in *Olfr151*- and in *Olfr16*-expressing neurons in their basal state. Dark and light green dots: genes expressed significantly higher in *Olfr15*- and in *Olfr16*-expressing neurons, respectively. (E) Distribution of the differentially expressed genes between *Olfr151*- and in *Olfr16*-expressing neurons in their basal state. (F) Scatter plot showing the differential expression analysis between *Olfr151-* and *Olfr16-* expressing neurons after exposure to their respective cognate ligands. Dark and light green dots: genes expressed significantly higher in *Olfr151*- and in *Olfr16*-expressing neurons after activation, respectively. (G) Volcano plot showing differentially expressed genes in *Olfr151*- and in *Olfr16*-expressing neurons after agonist exposure. Dark and light green dots: genes expressed significantly higher in *Olfr15*- and in *Olfr16*-expressing neurons, respectively. (H) Distribution of the differentially expressed genes between *Olfr151*- and in *Olfr16*-expressing neurons after agonist exposure. (I) Volcano plot showing differentially expressed genes in *Olfr151*- and in *Olfr16*-expressing neurons after exposure to their respective cognate ligands. Orange and blue dots: genes significantly upregulated and downregulated in both *Olfr151*- and *Olfr16*-expressing neurons, respectively, after agonist exposure. Light orange and blue dots represent genes modulated in either *Olfr151*- and in *Olfr16*-expressing neurons, respectively. (J) Gene Ontology analysis of the common differentially expressed genes in *Olfr151*- and in *Olfr16*- expressing neurons. The dashed line corresponds to the significant threshold (FDR adjusted *p*<0.05).

### Modulation of transcription following odorant exposure

Modulations in mRNA concentration may result from various processes. Among them and first in line, the regulation of transcriptional activity and the modulation of mRNA half-life. Our previous work has pointed to a very rapid downregulation of odorant receptor gene mRNA concentration following odor exposure (as fast as 20 minutes), and in some cases an almost absence of odorant receptor mRNA 5 hours after stimulation (von der Weid *et al*., 2015). This nearly immediate modulation and complete loss of messenger is suggestive, or at least compatible with an active degradation of cytosolic mRNAs. To explore this question, we took advantage of the different characteristics of nascent and mature mRNAs, namely the presence and lack of intronic sequences, respectively. We first analyzed the exonic versus intronic reads of the modulated genes that we identified after odorant exposure of Olfr151 and Olfr16 neurons (Figure 5A). Following agonist exposure, a large portion of the genes whose modulation was determined after analysis of exonic reads, were also modulated, both up and down and in both Olfr151- and Olfr16-expressing neurons, when restricting the analysis to intronic reads (Figure 5B-G). To further explore this question, we exposed wild type mice to ethyl isobutyrate (Figure 5H), an agonist for which we previously determined a set of highly responsive odorant receptor genes, among which *Olfr60*, *Olfr166* and *Olfr169*, whose corresponding mRNA concentrations drastically decrease after exposure. We evaluated this potential modulation at the level of nascent mRNAs. We observed a downregulation of odorant receptor mRNAs that was similar using exonic and intronic reads as readouts (Figure 5I). Finally, we looked at the cellular localization of transcripts (Figure 5H). We took advantage of the very high level of odorant receptor transcription that makes nascent transcripts easily visualized using in situ hybridization. Since odorant receptors are transcribed monoallelically, a single nuclear transcriptional spot corresponding to their expressed odorant receptor gene is observed in the nucleus of each sensory neuron (Figure 5J). We exposed wild type mice to ethyl isobutyrate and performed in situ hybridizations with a probe specific for *Olfr171*, whose mRNA we previously showed to be modulated following ethyl isobutyrate stimulation. We quantified the intensity of the nuclear signal, before and after agonist exposure, and found a significant decrease in signal intensity after ethyl isobutyrate exposure (Figure 5J). Taken together, these data all point to an odorant-induced modulation of transcription, and not to a mechanism involving degradation or stabilization of mature mRNAs.

**Figure 5.**
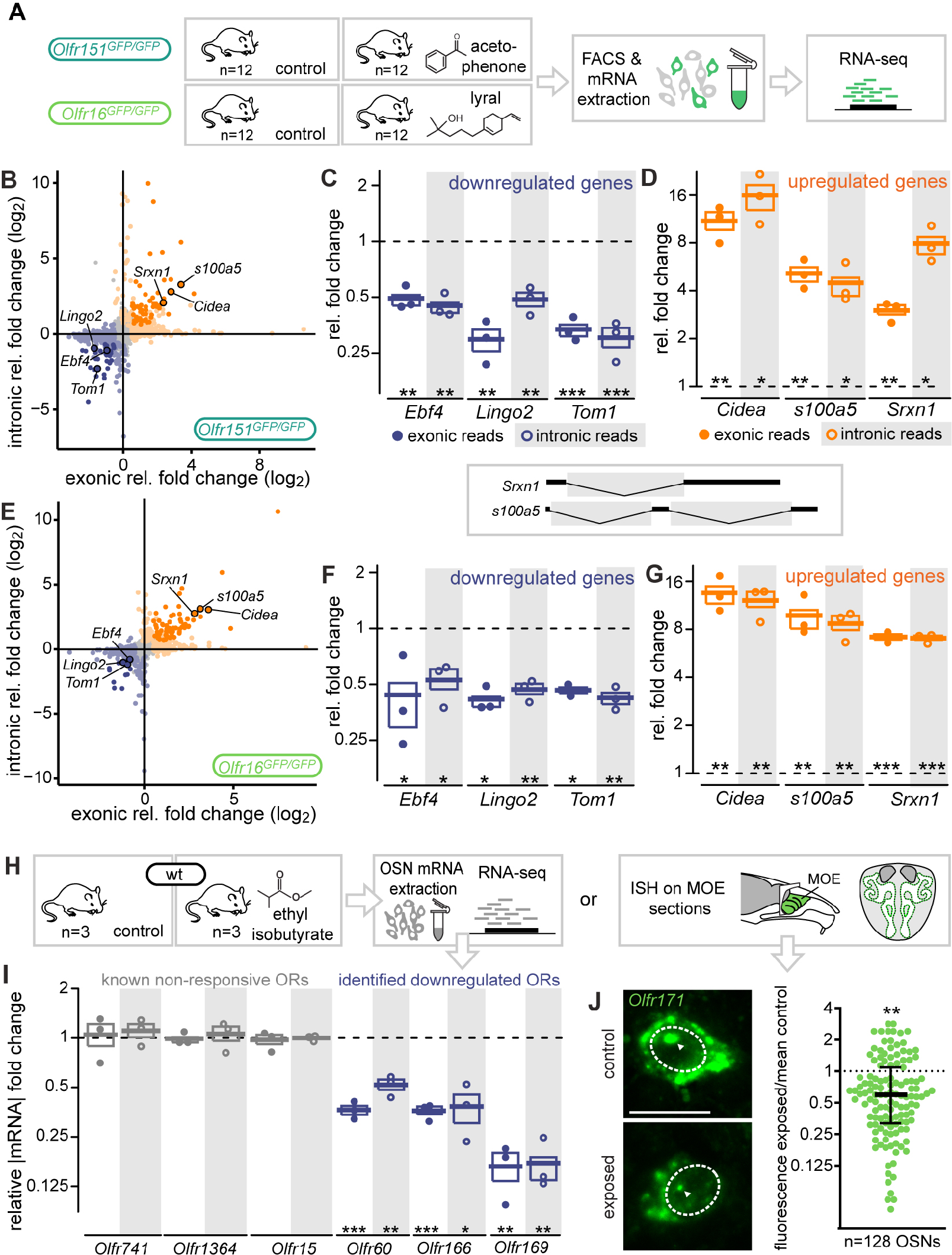
Odorant-induced modulations of mRNA levels result from transcriptional regulation. (A) Schematic of the experiment. After being exposed to their cognate ligand, fluorescent neurons from *Olfr151*^GFP/GFP^ and *Olfr16*^GFP/GFP^ mice were FAC-sorted and total mRNA was sequenced. (B) Scatter plot showing fold changes observed using exonic or intronic reads of FAC-sorted Olfr151- expressing neurons. Each gene is represented by a dot. (C) Examples of downregulated genes in *Olfr151*-expressing neurons (shown in (B)) after acetophenone exposure, whose fold change relative to control conditions were measured using exonic and intronic sequences. \*\**p*<0.01, \*\*\**p*<0.001, two sample independent t test using a linear model in R adjusted for FDR multiple comparisons. (D) Examples of upregulated genes in *Olfr151*-expressing neurons (shown in (B)) after lyral exposure, whose fold change relative to control conditions were measured using exonic and intronic sequences. \**p*<0.1, \*\**p*<0.01, two sample independent t test using a linear model in R adjusted for FDR multiple comparisons. (E) Scatter plot showing fold changes observed using exonic or intronic reads of FAC-sorted Olfr16- expressing neurons. Each gene is represented by a dot. (F) Examples of downregulated genes in *Olfr16*-expressing neurons (shown in (B)) after acetophenone exposure, whose fold change relative to control conditions were measured using exonic sequences. \**p*<0.1, \*\**p*<0.01, two sample independent t test using a linear model in R adjusted for FDR multiple comparisons. (G) Examples of upregulated genes in *Olfr16*-expressing neurons (shown in (B)) after lyral exposure, whose fold change relative to control conditions were measured using exonic and intronic sequences. \*\**p*<0.01, \*\*\**p*<0.001, two sample independent t test using a linear model in R adjusted for FDR multiple comparisons. (H) Schematic of the experiment where after exposure of wild type mice to ethyl isobutyrate, the olfactory transcriptome was either analyzed by RNA sequencing (I) or the main olfactory epithelium was sectioned and hybridized *in situ* with probes recognizing *Olfr171* transcripts (J and H). (I) The downregulation of three odorant receptor transcripts (corresponding to receptors activated by ethyl isobutyrate) was observed at the level of both intronic and exonic reads. \*\*\**p*<0.001, two sample independent t test using a linear model in R adjusted for FDR multiple comparisons. (J) Representative images of a control (top) and ethyl isobutyrate-exposed (bottom) olfactory sensory neuron, hybridized with an *Olfr171* probe. Dashed lines delimit the nucleus, and arrowheads point towards the site of transcription of *Olfr171*. Scale bar, 10 um. H) Fluorescence intensity within the transcription foci. Each dot represents an ethyl isobutyrate-exposed neuron. Fluorescence intensity was divided by the mean of all control neurons (median +/- 25th to 75th percentile). \*\**p*=0.0017, unpaired t test with Welch’s correction.

### Adenylyl-cyclase-dependent activity-induced transcriptomic adaptation

Following our identification of the nucleus as the source of the activity-dependent modulation of mRNA levels, we aimed at identifying the molecular events that underlie this phenomenon. We hypothesized that the canonical olfactory transduction cascade, which is well understood and involves multiple players, was likely involved (Figure 6B). We thus genetically dissected the cascade to evaluate the role that each element may play in the odorant-induced transcriptional response. We used several transgenic mice that were deficient in specific cascade elements (colored in Figure 6B), exposed these mice in vivo to ethyl isobutyrate for 5 hours, extracted their olfactory mRNAs, and used as a main readout the downregulation of the odorant receptor genes known to be affected by odorant exposure (Figure 6A,C). We evaluated the odorant-induced transcriptomic response of a null mutant of arrestin b2 (*Arrb2^del/del^*), a guanine nucleotide-binding protein *γ*8 subunit null mutant (*Gng8^del/del^*), an *Omp* null mutant (*Omp^del/del^*), a null mutant of the cyclic nucleotide gated channel subunit *α*4 (*Cnga4^del/del^)*, a conditional, olfactory-specific, null mutant of the cyclic nucleotide gated channel subunit *α*2 (*Omp^cre^;Cnga2 ^flox/Y^*), and a conditional, olfactory-specific null mutant of adenylyl cyclase 3 (*Gng8^cre^;Adcy3^flox/flox^*). To the exception of *Gng8^cre^;Adcy3^flox/flox^* mutants, none of the mutant lines bearing transduction cascade null alleles exhibited any deficiency in the downregulation of the known ethyl isobutyrate-responsive odorant receptor genes (Figure 6E-H). *Gng8^cre^;Adcy3^flox/flox^* mutants showed a complete absence of activity-induced transcriptional modulation of the genes of interest (Figure 6I). Since the approach involved whole tissue RNA extraction, the evaluation of the downregulated genes was limited to odorant receptor genes, the downregulation of other genes being invisible due to their expression in non-responsive neurons and their lack of downregulation in these latter (Figure S6A). We however expanded our analysis by evaluating the expression of genes (*Krsr2*, *Srxn1*, *Mustn1*, *S100a5*, *Cd24a* and *Epha5)* that we found strongly upregulated after agonist exposure of both Olfr151 and Olfr16-expressing neurons (Figure S6B). We reasoned that 1) these genes were also likely to be upregulated after ethyl isobutyrate exposure and 2) that despite the transcription of these genes in ethyl isobutyrate non-responsive neurons and thus a significant dilution of the upregulation (Figure S6A) (a dilution of over 120x if 10% of the sensory neurons respond to ethyl isobutyrate), we may still be able to detect some signal. According to our prediction, we found these genes upregulated in the olfactory epithelium of ethyl isobutyrate-exposed mice (Figure S6B). When the same protocol was applied to *Gng8^cre^;Adcy3^flox/flox^* mice, the upregulation was abolished (Figure S6B). Adenylyl cyclase 3 thus plays a critical role in the odorant-induced modulation, a modulation that does not involve the downstream cyclic-nucleotide gated channel.

**Figure 6.**
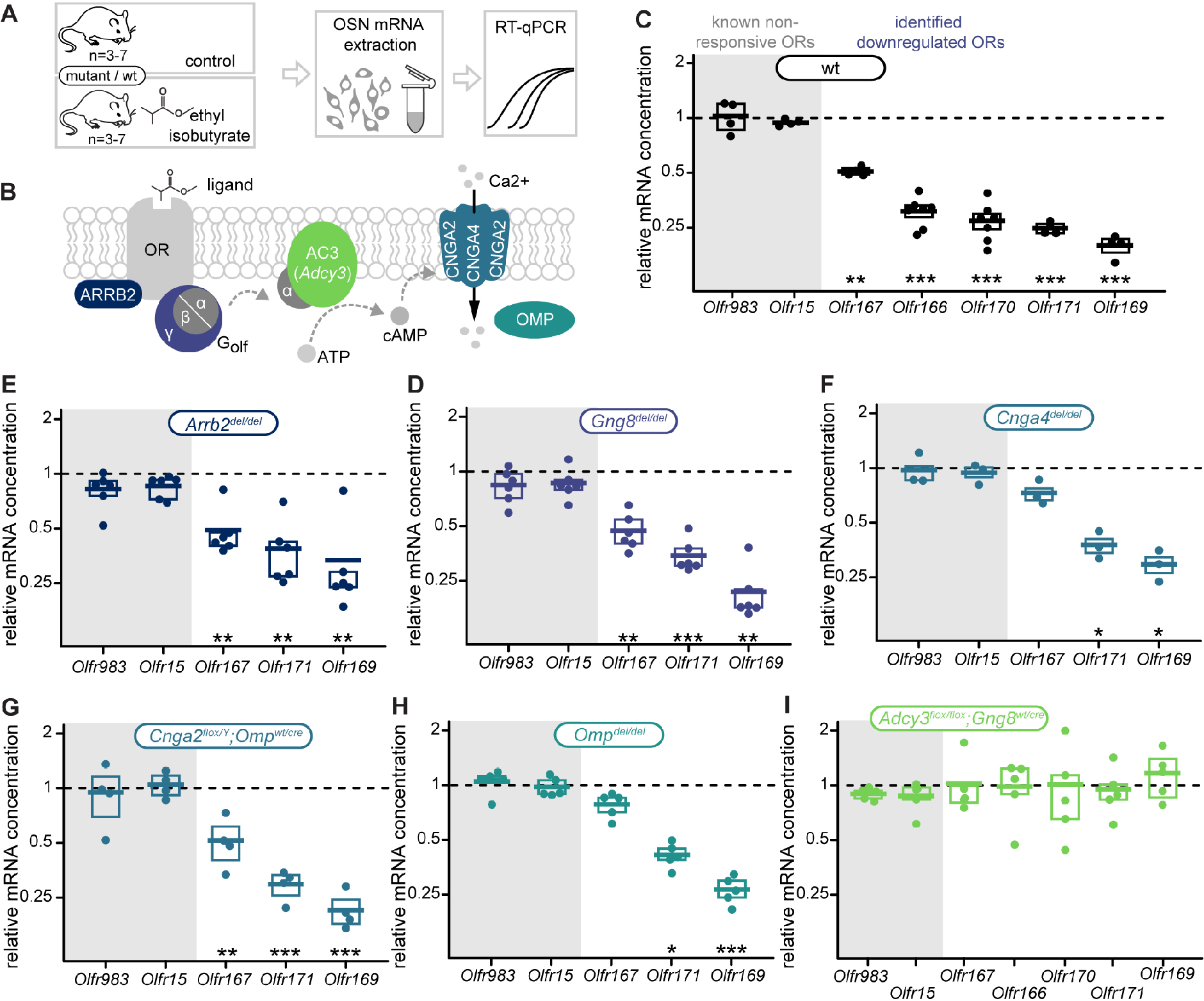
Adenylyl cyclase 3 as a critical player in the odorant-induced transcriptomic reprogramming. (A) Schematic of the experiment. Wild-type and mutant mice lacking different elements of the olfactory transduction cascade were exposed to ethyl isobutyrate. The modulation of gene expression levels of known non-responsive and responsive odorant receptor genes, and genes we found modulated after activation of both *Olfr151*- and *Olfr16*-expressing neurons, was evaluated by RT-qPCR. (B) Schematic of the olfactory transduction cascade. All elements highlighted in colors were individually targeted. (C) Transcriptional downregulation of odorant receptor genes in wild-type mice exposed to ethyl isobutyrate. *Olfr983* and *Olfr151* were known not to respond to this ligand, while *Olfr167*, *Olfr166* and *Olfr170* were previously shown to respond to this ligand. \**p*<0.1, \*\**p*<0.01, \*\*\**p*<0.001 (FDR adjusted), two sample independent t test using a linear model. (D-I) Transcriptional downregulation of OR genes in mutant mice lacking elements of the olfactory transduction cascade. \**p*<0.1, \*\**p*<0.01, \*\*\**p*<0.001 (FDR adjusted), two sample independent t test using a linear model.

## Discussion

We took advantage of the unique opportunity provided by the mouse olfactory epithelium, which contains hundreds of singular cell subpopulations, each defined by the expression of a known chemoreceptor gene and thus specifically activable at will in vivo, to explore the diverse signals that may determine and modulate neuronal transcriptional identities. Our results point to an unexpected tuning of cell identities in this sensory system. We found that the transcriptomic identity of mouse olfactory sensory neurons is remarkably variable, a variability that results from the interplay of two dimensions: a first, steady state transcriptomic identity that characterizes each population, to which a second may be added, driven by the recent activity of the sensory neuron.

How different are the hundreds of olfactory sensory neuron transcriptomes at rest? These transcriptomic dissimilarities can be divided into two categories. First, those that involve genes whose mRNA levels are relatively narrowly tuned and specific for any given population, providing a range of expression which is barely overlapping between populations. Then those that involve genes whose expression may be very high in some neurons, and completely absent in others. The combination of these two criteria generates a code, which is exclusive to every odorant-expressing population. Following odorant-mediated activation, we found a significant overlap between modulated genes. These represent late responsive genes (in opposition to early response genes). Late regulated genes typically encode proteins that regulate dendritic growth, synapse elimination or spine maturation (West and Greenberg, 2011). In our case, which possibly reflects the peculiarly ordered and relatively invariant olfactory circuitry, they mostly pertain to signal transduction categories. Interestingly, among the main genes involved in the olfactory transduction cascade, including those coding for odorant receptors, *Gna1*, *Adcy3*, *Cnga2*, *Cngb1b* or *Cnga4*, none were upregulated after odorant-mediated activation, and most were downregulated (data not shown). Such transcriptomic contraction affecting all members of the olfactory cascade suggests a functional transcriptional adaptation, leading to a decreased response to the experienced odor.

The expressed odorant receptor appears to be the main determinant in the establishment of the olfactory sensory population transcriptomic identities, both for steady-state and odorant-induced identities. Indeed, for this latter, the response profile of a neuron to agonists depends on the functional of the chemoreceptor it expresses, and thus on its sequence. Relative to steady state identity, our data show that transcriptomic similarities are better predicted by receptor similarities than by shared regulatory sequences represents a strong argument in favor of a direct role played by the olfactory receptor in basal transcriptomic identity.

But how to explain such critical role played by the odorant receptor in neurons at rest? Naturally, the resting state in a laboratory setting is an imperfect proxy of an odor void, a few sensory neurons always firing in a nose even without specific experimental stimulation, since some volatile molecules are always present in the environment. But without specific exposure to an odorant source, most olfactory sensory neurons are silent, thus preventing the establishment of a readable molecular dynamic range across neuronal populations that may accommodate for hundreds of specific cellular states (that are necessary to translate into hundreds of different transcriptional profiles). An interesting candidate mechanism for the generation of such odorant-receptor-dependent and odorant-independent graded cellular states is the agonist-independent natural basal activity of GPCRs. G-coupled receptors are indeed known to spontaneously oscillate between two conformations, one active and the other inactive, in the absence of ligands (in the presence of these latter, the receptors are stabilized in an active state (Kobilka and Deupi, 2007)). This effect was first observed, over 30 years ago, with the delta opioid receptor (Costa and Herz, 1989). Since then, and despite being often considered as noise, many more examples of such constitutive GPCR activity have been reported (with some receptors exhibiting very high levels of agonist-independent activity such as the Ghrelin receptor (Damian et al., 2012)), as well as various diseases associated with GPCR mutations affecting this constitutive activity. In the olfactory system, constitutive odorant receptor activity has been described and plays an important role: it regulates anterior-posterior targeting of olfactory sensory neurons during development (Nakashima et al., 2013). Constitutive odorant receptor activity thus appears to produce several different cellular states, whose number would be sufficient for defining enough non-overlapping neuronal categories to which distinctive rostro-caudal projection positions are assigned.

cAMP is a major player in olfaction, and the broad and evolutive transcriptional landscape we describe here appears to also involve this second messenger. cAMP function is pivotal at the level of the establishment of the system, as well as for its function, and is possibly directly involved in the transcriptional modulations we report here. First, cAMP plays a role in the process that leads to restricting odorant receptor genes to a single gene and allele (Dalton et al., 2013; Lyons et al., 2013) (although its critical role in this exclusion mechanism is debated (Movahedi et al., 2016)), a process involving a negative feedback signal resulting from the expression of the chosen receptor gene (Lewcock and Reed, 2004; Serizawa et al., 2003). Second, cAMP is essential to the establishment of a spatial map in the olfactory bulb, that requires, late during olfactory neuron maturation, the expression of glomerular segregation molecules such as Kirrel 2 and 3 (Nakashima *et al*., 2013; Serizawa et al., 2006). This process used the canonical olfactory transduction cascade, which is somehow activated during development. This cascade involves, after receptor activation, the release of G*α*olf subunits which activate adenylyl cyclase 3, whose product (cAMP) then facilitates the opening of multimeric cAMP-activated Na^+^/Ca^++^ ion channels composed of CNGA2, CNGA4 and CNGB1b. The following influx of calcium ions leads to the opening of calcium-activated chloride channels, to the efflux of Cl^-^ from the neurons, and to action potentials. Third, via this very cascade, cAMP is part of the odorant-induced signal transduction in mature neurons. The cascade involves, after receptor activation, the release of Gαolf subunits which activate adenylyl cyclase 3, whose product (cAMP) then facilitates the opening of multimeric cAMP-activated Na^+^/Ca^++^ ion channels composed of CNGA2, CNGA4 and CNGB1b. The following influx of calcium ions leads to the opening of calcium-activated chloride channels, to the efflux of Cl^-^ from the neurons, and to action potentials. This olfactory cascade is possibly involved in the activity-driven transcriptional reprogramming we report here, since *Adcy3* null mice are immune to the odorant-induced transcriptomic adaptation. However, if true, given the maintenance of the odorant-induced transcriptional responses in CNG channel mutants, the following steps of the canonical cascade are not part of the transcriptional modulation process. In fact, action potentials are not even required for agonist-dependent transcriptional modulation, *Cnga2^del/del^* olfactory neurons being silent. A last and independent role of cAMP is played earlier, during olfactory circuit formation, at the time of the targeting of olfactory sensory neurons to the olfactory bulb (Chesler et al., 2007; Dal Col et al., 2007; Imai et al., 2006; Nakashima *et al*., 2013; Zou et al., 2007). During neuronal maturation, different levels of cAMP are produced in different immature sensory neuron populations, which translate in the differential expression of guidance genes, in particular *Nrp1* (Dal Col *et al*., 2007; Imai *et al*., 2006). These different levels result from the basal activity of olfactory odorant receptors we previously discussed (Nakashima *et al*., 2013).

To further explore the molecular players involved in the cAMP-dependent transcriptomic modulation, some potential targets of cAMP appear obvious. In particular the cAMP response element-binding protein (CREB), a transcription factor critical for activity-dependent neuronal plasticity (West and Greenberg, 2011) and involved in the activation-induced prolonged lifespan of olfactory sensory neurons (Watt et al., 2004) (the evaluation of the role played by this factor may however prove difficult to explore, due to the known redundancy of other members of the CREB family). Interestingly, a histone H2B variant termed Hist2H2BE exhibits levels of expression in olfactory sensory neurons that are associated with the identity (defined by the expressed odorant receptors) of the different neuronal populations. Its expression modulates neuronal longevity, is reduced after sensory activity, and possibly participates in activity-dependent changes (Santoro and Dulac, 2012). This downregulation of Hist2H2BE is dependent on adenylyl cyclase 3 (Ac3) function, which again, means on cAMP.

The mammalian olfactory mucosa thus represents a multifunctional sensor whose neuronal elements, that use olfactory receptors as internal and external probes, are in constant evolution, adapting to the world via the activation of large-scale transcriptomic programs. Whether this extreme transcriptomic diversity and dynamics functionally parallels the one recently observed in some central circuits (Saunders et al., 2018; Tasic et al., 2016; Tasic et al., 2018; Zeisel et al., 2018), and whether the molecular tools involved in transcriptomic adjustments are shared between different circuits, remains to be explored. Given the extraordinary transcriptomic diversity of neurons in the mammalian brain and knowing that 90% of non-sensory GPCRs are expressed in mammalian brains, the question is worth asking.

## Acknowledgments

We thank Chenda Kan and Véronique Pauli Jungo for expert technical assistance, and the iGE3 Genomics Platform at the University of Geneva for assistance during bulk and single cell RNA-seq experiments. This research was supported by the University of Geneva and the Swiss National Science Foundation (grant numbers: 31003A_172878 to A.C. and 310030_189153 to I.R.).

## Author contributions

IR and AC conceived the project, acquired funding, supervised research, and wrote the manuscript. LFH, AA, LF, LM, and AH participated to the experimental design, performed experiments, collected data and interpreted results. MB participated to the interpretation of the results and visualization of the data. JT participated to the experimental design, interpretation of the results, visualization of the data, and performed the analysis of scRNAseq data.

## Data and code availability

Bulk and single-cell RNA-seq data have been deposited in NCBI GEO and can be accessed using the superseries GSE185168 accession number, and are publicly available as of the date of publication. All original code used to analyze data reported in the paper are provided at the GitHub repository https://github.com/irlabgenev/DREAM_pt2

## Experimental model and subject details

### Animals

C57BL/6J male mice were purchased at 5-7 weeks of age from Charles River Laboratories. Upon arrival, they were housed in groups of 4-5 animals in standard type II cages with access to food and water *ad libitum*. The following transgenic mouse lines were employed: *Adcy3^flox^, Cnga2^flox^, Gng8^cre^, Omp^cre^, Omp^GFP^, MOR23(Olfr16)^GFP^, M71(Olfr151)^GFP^* (Assens et al., 2016; Li et al., 2004; Matsuo et al., 2015; Potter et al., 2001; Skarnes et al., 2011; Vassalli et al., 2002). The *Cnga4^del^* allele was generated by homologous recombination of a targeting vector lacking the sequences encoding for the 3^rd^ and 4^th^ transmembrane domains, the channel pore, the cyclic nucleotide-binding regions and the intervening introns 3, 4 and 5. The resulting deletion is 2528bp long, and starts with an AvrII site in intron 3 and ends with a XhoI site in exon 6. Routine experiments were performed in a room with a 12 hours light-dark cycle, the light phase lasting from 6:30 a.m. to 6:30 p.m. All animals were housed and treated in accordance with the veterinary guidelines and regulations of the University and of the state of Geneva.

### Method details

#### 10X single-cell RNA sequencing

##### Cell isolation, sorting and sequencing

8-weeks-old male C57BL/6J mice were used (n=4). All experiments were performed during daytime. Mice were euthanized with intraperitoneal injection of pentobarbital (150 mg/Kg) and their olfactory epithelia were immediately extracted and processed for tissue dissociation using the Papain Dissociation System (cat #LK003150; Worthington® Biochemical Corporation, New Jersey, USA) following the manufacturer’s protocol. Cell suspensions were then incubated with 2 μg/ml of Hoechst 33342 (a UV fluorescent adenine-thymine binding dye; #H1399, Life Technologies) at 37°C for 15 min. Before fluorescence activated cell sorting (FACS) and to exclude dead cells, 1 μM of DRAQ7^TM^ (a far-red fluorescent DNA intercalating dye; #DR71000, BioStatus) was added to the cell suspensions. Approximately 80’000 Hoechst+/DRAQ7-cells were collected from each sample, each in a final volume of 100 μl. After FACS sorting, cell suspensions were concentrated at 800 cells/μl. The targeted cell recovery was set to 10’000. In accordance with the Cell Suspension Volume Calculator Table of 10X Genomics, 22.6 μl of nuclease-free water was added to 20.6 μl of cell suspension and the samples were loaded on the 10X Genomics Chromium controller. GEM generation and barcoding, cDNA amplification and cDNA library construction were performed following the 10X Genomics Chromium Next GEM Single Cell 3’ v3.1 protocol (dual index libraries). The cDNA libraries from each sample were then pooled and loaded at 2 nM on 2 lanes of the Illumina HiSeq 4000 system for paired-end sequencing.

##### scRNA-seq mapping and counting

fastq files were pre-processed with Cell Ranger version 6.0.1 (Zheng et al., 2017) with default settings. Reads were mapped on the *Mus musculus* genome primary assembly reference 38 (GRCm38) using the STAR aligner (Dobin et al., 2013) implemented in Cell Ranger. A modified version of the Ensembl release 102 of the *Mus musculus* GTF annotation was used. This GTF file was updated with the re-annotation of the 3’UTR of olfactory receptor genes. The filtered feature-barcode matrices were used for downstream analysis. These matrices included a total of 21,809 cells (sample 1: 5,364 cells; sample 2: 5,696 cells; sample 3: 4,756 cells; sample 4: 6,004 cells).

##### scRNA-seq data filtering

Single-cell RNA sequencing data analyses were performed on R version 4.0.5 using the Seurat R package version 4.0.1 (Butler et al., 2018; Satija et al., 2015). Seurat’s functions were used with default settings unless specified. The standard analysis consisted of the following steps. First, the four 10X gene expression matrix files were individually loaded into R using the *Read10x* function of Seurat. The 10X data were then converted to Seurat objects using the *CreateSeuratObject* function of Seurat. The gene expression data was then normalized using the *SCTransform* function of Seurat (Hafemeister and Satija, 2019), and the top 5,000 variable genes were determined for datasets integration (Hao et al., 2021; Stuart et al., 2019). Following the integration of the four datasets, a preliminary clustering was performed without any additional cell filtering in order to identify and remove cell clusters composed of blood, immune or suffering cells (i.e. cells exhibiting high expression levels of mitochondrial genes). Principal component analysis (PCA) was performed on the integrated assay of the Seurat object using the *RunPCA* function of Seurat. A visual inspection of their explained standard deviation led to the selection of the top 9 PCs for subsequent cell clustering. To construct a shared nearest-neighbor graph, the above-mentioned PCs were used as input to the *FindNeighbors* function of Seurat (dims = 1:9). Cell clusters were then identified using the *FindClusters* function of Seurat with a clustering resolution of 1. This preliminary clustering yielded 27 cell clusters. Cluster-specific gene markers were then identified for cluster annotation. Briefly, the raw dataset containing cells sampled from all four mice was normalized by library size, scaled to 10^4^ and natural-log-transformed after adding a pseudocount of 1 using the *NormalizeData* function of Seurat. This normalized data was then used for differential expression analysis computed between each cell cluster and all other clusters taken together using the Wilcoxon rank sum test implemented in the *FindAllMarkers* function of Seurat (test.use = “wilcox”; only.pos = TRUE). Only genes with an adjusted p-value below 0.05 were considered. A blood cell cluster (n = 1) was identified based on its high expression of hemoglobin chain complex genes such as *Hba-a1* and *Hba-a2*. Immune cell clusters (n = 7) were identified based on their high expression of known immune cells markers such as *Igkc*, *Cd52*, *Cybb*, *Ctss* and *Tyrobp*. Suffering cell clusters (n = 4) were identified based on their high percentage of mitochondrial gene counts. This preliminary clustering led to the removal of 12 cell clusters from the dataset (n = 4,376 cells). Furthermore, cells were also filtered out if their percentage of mitochondrial counts exceeded 10% of their total counts or if they expressed less than 1,000 genes (n = 1,506 cells). This preliminary analysis resulted in retaining 15,927 cells.

##### scRNA-seq clustering and analysis of main olfactory epithelium cells

The retained cells were used to identify cell clusters composing the mouse main olfactory epithelium (MOE). The corresponding dataset was normalized and integrated as described in the previous paragraph (see *scRNA-seq data filtering*) with the following differences: the first 15 PCs were used for the *FindNeighbors* function of Seurat and a resolution of 0.3 was used for the *FindClusters* function of Seurat. This analysis led to the identification of 13 cell clusters. Cluster identities were then determined from the differentially expressed genes in each cluster (see above for more details). The markers described in Fletcher et al. 2017 (Fletcher *et al*., 2017) were used for the annotation of the mouse MOE cell types. From the 13 clusters, 5 corresponded to mature OSNs (mOSNs) based on their high expression of *Omp*, *Cnga2* and *Gng13* but not *Gap43* (Figure S1A) These clusters were then merged together into only one cluster of mOSN (Figure 1B). To visualize the resulting 9 cell clusters on a 2-dimensional plot, the uniform manifold approximation and projection (UMAP) (Becht et al., 2018; McInnes et al., 2018) plot was computed using the *RunUMAP* function of Seurat and the first 15 PCs previously selected (dims = 1:15) (Figure 1B).

Mature OSNs were selected from the main olfactory epithelium dataset for downstream analyses (n = 10,737 cells). For each mOSN, the detected olfactory receptors were ordered based on their expression levels: 7,178 OSNs displayed the expression of a single olfactory receptor, 2,701 OSNs displayed the expression of two olfactory receptors and 702 OSNs displayed the expression of at least three olfactory receptors. To remove cells that could correspond to multiplets (among those co-expressing multiple olfactory receptors), the distribution of the expression levels of the highest expressed receptors was analysed using the log normalized data. An “is expressed” cutoff was set at three median absolute deviations from the median of the levels of expression of the highest expressed receptors. OSNs whose highest expressed receptor had an expression level below this cutoff were removed from the dataset (n = 223 cells). Moreover, OSNs that expressed more than one receptor at an expression level higher than this cutoff were also filtered out from the dataset (n = 358 cells). Finally, roughly 1.5% of the mOSNs (n = 156 cells) did not show receptor expression and were also discarded from the dataset. The OSN population identity of each of the remaining cells (n = 10,006 cells) was then determined based on the olfactory receptor that displayed the highest expression level in that given cell. This led to the identification of 955 odorant receptor (OR)-expressing OSN populations (n = 9,959 cells) and 7 TAAR-expressing OSN populations (n = 44 cells), as well as a *Gucy1b2*-expressing OSN population (n = 3 cells). OSN populations represented by at least 3 cells in the dataset were included for clustering and downstream analyses (n = 9762 cells).

Two parallel analyses were carried out: by keeping or removing the olfactory receptor genes from the count matrix. The corresponding datasets were normalized and integrated as described above (see *scRNA-seq data filtering*) with the following differences: the percentage of mitochondrial gene counts were used as confounder variables in the *SCTransform* function of Seurat (vars.to.regress = “percent.mt”); the first 18 or 19 PCs were used for the *FindNeighbors* and *RunUMAP* functions of Seurat for the analyses including or not the olfactory receptor genes, respectively; and a resolution of 1.9 or 1.7 was used for the *FindClusters* function of Seurat for the analyses including or not the olfactory receptor genes, respectively. These concurrent analyses led to the identification of 25 (including olfactory receptor genes) or 23 (not including olfactory receptor genes) cell clusters, respectively. Cluster identities were then determined from the differentially expressed genes in each cluster (see *Data filtering* for more details). After cluster merging, a total of 10 clusters were retained, which were then subdivided into groups of “dorsal” or “ventral” clusters based on their complementary expression of *Nqo1* (a dorsal mOSN gene marker) or *Nfix* (a ventral mOSN gene marker), respectively. These broad clusters were each composed of five sub-clusters characterized by their expression of specific markers genes or absence of them: Dlg2+, Calb2+, Cd55+, Cd36+ and Dlg2-;Calb2-;Cd55-;Cd36- clusters. The clustering similarity between the two analyses (i.e. including or excluding the olfactory receptor genes) was computed with the normalized mutual information metric using the *compare* function of the igraph R package version 1.2.6 (method = “nmi”).

The pairwise transcriptomic Euclidean distances between pairs of OSNs was computed on the first 19 PCs of the mOSN dataset (computed from the sctransform-normalized and integrated count matrix; see *Data filtering* for more details). Euclidean distances were then split into two categories: distances between pairs of OSNs expressing the same olfactory receptor (intra) or pairs expressing different receptors (inter). In order to test if the difference in the distributions of Euclidean distances between pairs OSNs expressing the same or different olfactory receptors was not a random effect, we permutated the cell identities (and hence the corresponding olfactory receptor identities) of the PCs prior to distance calculation (n = 1000 permutations). The Wilcoxon rank sum test was then used to compare these distributions and the p-values were adjusted for multiple comparisons using the Bonferroni correction method.

Similar to what was performed per cluster, OSN population-specific gene markers were identified using the log normalized UMI counts and the *FindAllMarkers* function of Seurat (test.use = “wilcox”; only.pos = TRUE). In Figure 1G, the largest OSN population from each cluster was selected and the gene expression levels of its cells were compared to those of all other cells from the dataset. In Figure 1H, the six largest OSN populations from the Dlg2-, Calb2-, Cd36-, Cd55- ventral cluster were selected and for each of these populations the gene expression levels of their cells were compared to those of all other cells from that specific cluster. Only genes expressed in at least 70% of the cells of the given population and that yielded an adjusted p-value below 0.05 were considered. For plotting, the log normalized data was scaled and centered using the *ScaleData* function of Seurat, and the extreme values were clipped and set to the lower and upper limit values of the 95% confidence interval of the data using the *clip.data* function of the fsbrain R package version 0.4.3 (lower = 0.025; upper = 0.975).

#### Transcriptomic, genomic and amino acid distances

##### Functional OR gene identification and OR phylogeny

The functional OR phylogeny was partly built from the same sequence set as used in (von der Weid *et al*., 2015). To constitute this set, OR coding sequences were identified de novo in the mouse genome assembly GRCm38 using TBLASTN searches with previously annotated mouse OR protein sequences as queries. The hits were manually curated to filter out putative non-functional receptors. The criteria to consider an OR to be functional was the conservation of evolutionary constrained residues (Niimura, 2013), the integrity of the seven transmembrane domains and the absence of intron within the coding sequence (de March et al., 2015), resulting in a set of 1141 putatively functional OR. After this filtering, 11 filtered out ORs were retrieved as they were found to be expressed in a monoallelic fashion in one or more OSNs, in our scRNA-seq data. For these ORs, we used coding sequences as annotated in Ensembl version 102. Notably, 8 of these 11 ORs have their coding sequence spanning two exons, with most of the coding sequence (covering the seven transmembrane domains) included in the last exon.

A multiple sequence alignment including the resulting OR protein sequence set was obtained with Clustal Omega v1.2.4 (Sievers et al., 2011), using the *--full* and *--full-iter* options. The resulting alignment was trimmed to keep the sites between the most conserved start methionines and the last position with less than 90% of gaps.

The maximum likelihood phylogeny of the mouse functional ORs was calculated with Phyml version 20120412 (Guindon et al., 2010) using the following parameters: -d aa -m JTT -f e -v e -c 4 -a e -s BEST -o tlr. The resulting tree was rooted on the node at the origin of class I and class II ORs.

##### Transcriptomic identity of mature OSN populations and pairwise distance metrics

OSN populations are referred to as OSNs expressing the same Olfr genes in a monogenic manner. Each OSN population was assigned to the transcriptomic cluster to which the majority of cells belong to. In case of equivalences, we assigned the transcriptomic cluster randomly. In Figure 2A, OSN populations with three or more cells are displayed around the phylogeny, whereas in Figure 2C and on, we kept OSN populations represented by 10 or more OSNs to reduce noise in pairwise distance statistics. Transcriptomic identities of these populations were defined as the centroids of the population transcriptomes in the same PCA that was used for the transcriptome clustering including as well the top 19 PCs. Pairwise transcriptomic distances between OSN populations were obtained by calculating the Euclidian distance between their respective centroids.

Pairwise genomic distance between Olfr genes was measured as the distance in base pairs between start codons of Olfr genes. For the 8 Olfr genes that have their start codon on another exon, we instead used the first position of the last coding exon. Genomic distances were only obtained between genes in the same chromosome.

Pairwise distances between adjacent genes were used to aggregate Olfr genes in cluster. For this, the sorted distances were split into two groups using the Jenks natural break optimization for k=3. In that manner, the middle break is used to separate unbiasedly two categories of distances: the smaller distances representing the intracluster distances and the longer distances representing the intercluster distances. Next, we calculated the mean and the standard deviation of the intracluster distances and defined the clustering threshold as the mean plus 3 times the standard deviation. Finally, gene clusters were obtained by aggregating neighboring genes whose genomic distance was closer to each other than the clustering threshold.

Pairwise amino acid difference was measured on the protein alignment that was used for the phylogenetic reconstruction. For a given pair of aligned sequences, each substitution was scored according to the Miyata amino acid replacement matrix (Miyata et al., 1979). Insertions were scored as the mean replacement scores of each additional amino acid. The sum of these scores gave the pairwise amino acid difference.

In Figure 2F, we defined thresholds of genomic distance and amino acid difference to attribute pairs of OSN populations as being close in terms of genomic proximity between the Olfr genes they express or in terms of sequence identity between their respective OR. For genomic proximity, we evaluated all intergenic distances between neighboring ORs belonging to the same cluster and chose the 95th percentile of this distribution as the threshold value for a pair to be considered close. For sequence identity, we evaluated all pairwise amino acid differences between ORs belonging to the same class and chose the 5th percentile of this distribution as the threshold value for a pair to be considered close. For Figure 2G, a pair was considered distant in terms of genomic proximity when the corresponding Olfr genes were located in different Olfr gene cluster. A pair was considered distant in terms of sequence identity when the amino acid difference between their corresponding ORs was higher than the threshold used to identify the close pairs.

#### Chemicals

Odorants were directly purchased from Sigma-Aldrich, ethyl isobutyrate (W242802), acetophenone (42163). Lyral was obtained as a generous gift from Dr. Christian Margot (Firmenich).

#### Odorant exposure

For all odorant exposures, on the day preceding the odorant exposure mice were isolated and single-housed in a standard type II long cage. Exposure assays started at 8:00 a.m. and lasted 5 hours. For the exposed condition, a cotton swab was imbibed with 200µL of 5% odorant in a DMSO solution and was placed in the cage, while for the control condition, a cotton swab was imbibed with 200µL of DMSO only and was placed in the cage.

#### FACS-seq

##### Cell isolation, sorting and sequencing

*Olfr16^GFP/GFP^* and *Olfr151^GFP/GFP^ mice*, corresponding to mice carrying the *_Olfr16_irestauGFP/irestauGFP* _(*Olfr16*_*tm2Mom*_) and *Olfr151*_*irestauGFP/irestauGFP* _(*Olfr151*_*tm26Mom*_)_ alleles respectively (Potter *et al*., 2001; Vassalli *et al*., 2002), were used to isolate single fluorescent OSN populations from the whole MOE. 7-week-old *Olfr16^GFP/GFP^* and *Olfr151^GFP/GFP^* were exposed as described above to lyral and acetophenone, respectively. Control mice from each transgenic line were exposed to DMSO only. For each condition, there were 3 samples, where each sample was constituted by a pool of 4 mice. After odorant exposure, mice were euthanized by intraperitoneal injection of pentobarbital (150mg/Kg), the whole MOE was extracted and OSNs were dissociated by adapting the protocol described in Kaur et al., 2013 (Kaur et al., 2013). Briefly, the collected epithelia were minced inside a tube containing a dissociation buffer (D-csyteine-HCl 1M, EDTA 100mM, Papain 0.3U/µL, DNAse I (Ambion) 2U/µL and DNAse I 10x buffer (Ambion), dissolved in freshly prepared and oxygenated cold aCSF). The aCSF composition was the following: 118mM NaCl, 25mM NahCO3, 10mM D-glucose, 2mM KCl, 2mM MgCl2, 1.2mM NaH2PO4, 2mM CaCl2. Samples were then placed at 37°C for a total of 25 minutes allowing enzymatic dissociation of the tissues, during which they were subjected to a trituration step every 5 minutes using polished glass pipettes. At the end of the dissociation, each sample was filtered through a 20µm Nylon filter (Falcon), and centrifuged for 5 minutes at 200G. The supernatant was discarded and replaced with ice-cold aCSF. Before FAC-sorting, samples were incubated at 37°C for 20 minutes with Hoechst 33342 (1mg/mL) to label live cells. Cell-sorting was performed on an AriaII (BD Biosciences) cell-sorter, gated on Hoechst and GFP fluorescence. Cells were collected directly in lysis buffer from the Qiagen RNeasy plus micro kit. For the *Olfr16^GFP/GFP^* mice, 100 cells were collected per individual, resulting in 400 cells per biological pool. For the *Olfr151^GFP/GFP^* mice, 50 cells were collected per individual, resulting in 200 cells per biological pool. The difference in the total number of cells collected per experiment derives from the original respective OSN population sizes in the epithelium (Bressel et al., 2016). The RNA extraction was performed according to the Qiagen RNeasy plus micro kit protocol. The SMARTer™ Ultra Low RNA kit from Clontech was used for reverse transcription and cDNA amplification (12 PCR cycles) according to the manufacturer’s specifications, starting with a total volume of 9.5 µL per sample as total RNA input. 200 pg of cDNA were used for library preparation using the Nextera XT kit from Illumina. Library molarity and quality was assessed with the Qubit and Tapestation using a DNA High sensitivity chip (Agilent Technologies). Libraries were pooled at equimolarity and loaded at 11 pM for clustering on a Single-read Illumina Flow cell for the *Olfr16^GFP/GFP^* experiment. Reads of 50 bases were generated using the SBS HS v3 chemistry on an Illumina HiSeq 2500 sequencer. Deep sequencing of the Olfr16 dataset yielded a mean of 37.2M (*±* 1.3M) short single-reads for the control condition, and a mean of 35.4M (*±* 4M) short single-reads for the exposed condition. For the Olfr151 experiment, libraries were loaded at 2 nM for clustering on an Illumina HiSeq 4000 sequencer. Deep sequencing of the Olfr151 dataset yielded a mean of 58.5M (*±* 1.5M) short single-reads for the control condition, and a mean of 62.1M (*±* 1.3M) short single-reads for the exposed condition.

##### FACS-seq mapping and counting

STAR (v.2.7.0, Dobin et al., 2013) was used to map the generated reads on the Ensembl *Mus musculus* genome primary assembly reference 38 (GRCm38) that included the IRES-tau-GFP sequence. Gene expression quantification was carried out using featureCounts version 1.6.3 (Liao et al., 2014).

##### FACS-seq data filtering

To filter out lowly- and non-expressed genes for each OSN population (Olfr16 and Olfr151), a count threshold was determined to exclude all genes with expression values below this threshold across either the 3 control or 3 exposed samples. Briefly, the density distribution of gene counts was used to calculate the local minimum and this value was set as the threshold.

##### FACS-seq gene expression analysis

The DESeq2 package (v.1.30.1) was then used to perform differential expression analysis. After fitting a negative binomial generalized linear model (GLM), the Wald test (two-tailed) was used to test for significance of gene expression at a log_2_ fold change threshold of 0.5. To control the false discovery rate, the Wald test p-values were adjusted for multiple comparisons using the Benjamini-Hochberg procedure (Benjamini and Hochberg, 1995).

##### Gene Ontology enrichment analyses

All Gene Ontology (GO) enrichment analyses were performed testing GO terms mapped to the differentially expressed genes (DEGs) common to both analyzed OSN populations (Olfr16 and Olfr151) against a background of GO terms mapped to all other genes commonly expressed in both OSN populations. DEGs were analyzed for Gene Ontology (GO) enrichment by the topGO package using the runTest function with the “classic” algorithm and the Fischer statistics. To control the false discovery rate, the p-values were adjusted for multiple comparisons using the Benjamini-Hochberg procedure (Benjamini and Hochberg, 1995). The result of the GO terms analysis were then plotted with the ggplot2 function in R.

#### RNA extraction for bulk RNA sequencing and RT-qPCR

After odorant exposure, mice were euthanized by intraperitoneal injection of pentobarbital (150mg/Kg). From each animal, total olfactory epithelia (lateral and septal) from each side were collected and transferred to a tube containing 500 µL of ice-cold lysis buffer (Quickgene RNA tissue kit SII), 5 µL of β-mercaptoethanol and an RNAse free stainless steel bead measuring 0.5 cm in diameter. Samples were homogenized for 30 s at 6 ms^-1^ with a FastPrep-24 instrument (MP Biomedicals). After homogenization, samples were either flash frozen at -80°C for future RNA extraction or directly processed. For RNA extraction prior to bulk RNA-seq and RT-qPCR, we followed the manufacturer’s protocol from the Kurabo QuickGene RNA tissue kit SII. At the final elution step, 50 µL of RNAse-free water was added to each column. Samples were then treated with DNAse as per the Life Technologies Ambion I DNAse kit protocol and stored at -80°C.For FACS-seq and single-cell RNA-seq, RNA extraction protocols are described in the corresponding sections.

#### Bulk RNA-seq

##### Sequencing

Mice were exposed to ethyl isobutyrate as described above. After RNA extraction, cDNA libraries were generated with the Truseq RNA and DNA sample preparation kits after selection of polyA-containing mRNAs. Adapters for RNA-seq multiplexing were added to the cDNAs. The cDNA libraries were sequenced with a HiSeq2500 Sequencing system, where 100-bp reads were generated. The raw data generated in this experiment was previously published in Von der Weid et al. (von der Weid *et al*., 2015).

##### Bulk RNA-seq mapping and counting

The mapping and counting of bulk RNA-seq data was performed exactly as described for FACS-seq data above (*FACS-seq mapping and counting*, *data filtering* and *expression analysis*).

#### RT-qPCRs

##### RNA quantitation

Wild-type and mutant mice were exposed to ethyl isobutyrate and RNA was extracted as described above. RNA quantitation was performed using a Nanodrop, and the RNA quality was assessed with the Agilent Technologies 2100 Bioanalyzer. The reverse transcription was performed with the Takara PrimeScript RT Reagent kit (RR037A) with 500 ng of starting RNA in a total volume of 10 µL. Specific primers were designed to amplify exon-exon junctions (if feasible) and to generate a 80-150 bp amplicon (Table 3). Each qPCR reaction, was run in technical triplicates for each sample and took place in a total volume of 10 µL composed of 2.5 µL of cDNA at 4 ng/µL and 7.5 µL of PowerUp SYBR Green master mix from Thermofisher® together with primers at 300 nM. A QuantStudio5 machine with the following qPCR program parameters was used: 2 minutes at 50°C, 10 minutes at 95°C, 40 cycles each lasting for 15 s at 95°C and 1 minute at 60°C.

**Table 3.**
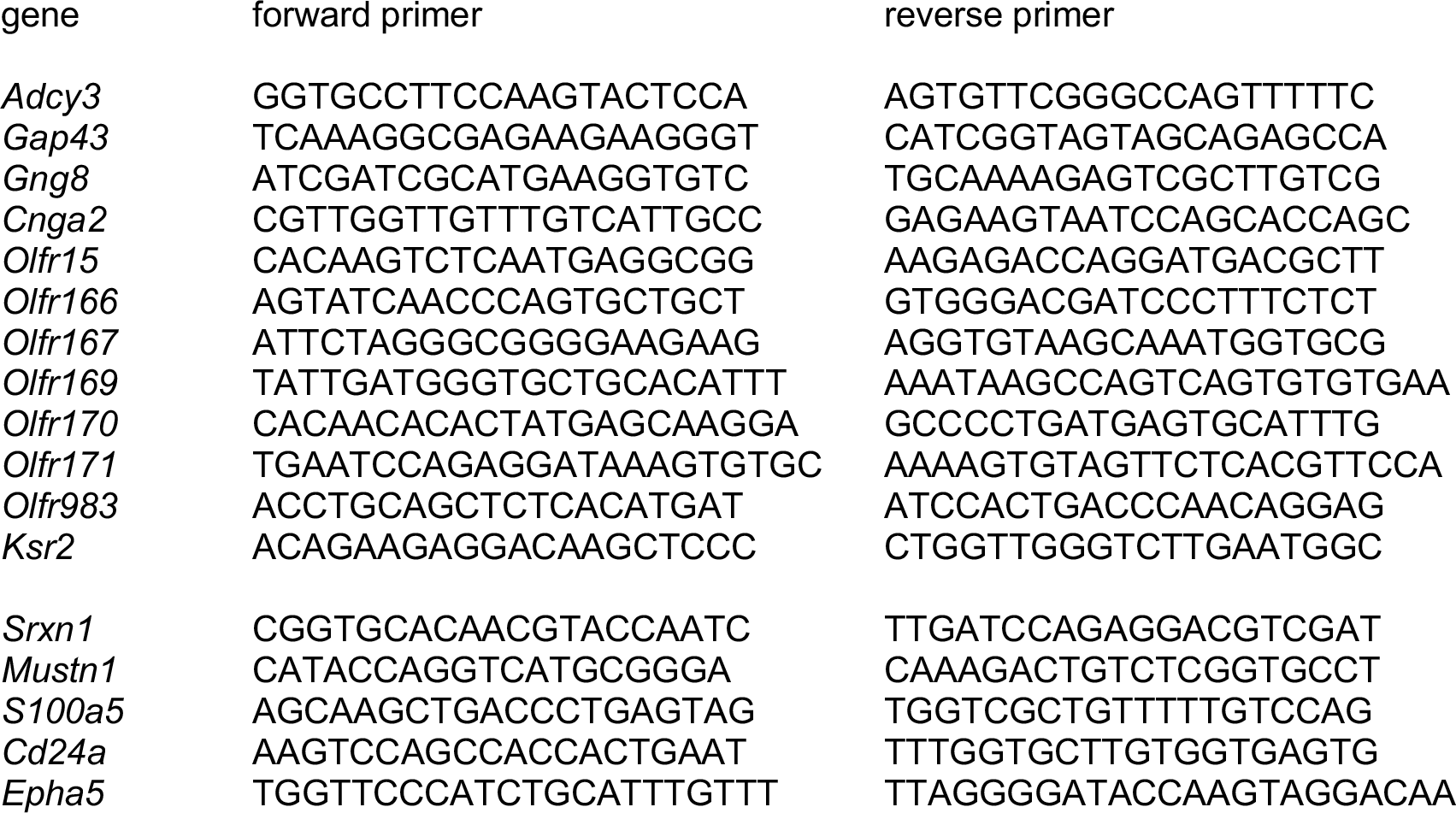
Primers used for RT-qPCR analysis.

##### RT-qPCR data analysis

Raw data were then analyzed using the ThermoFisher Cloud algorithm, setting manually a detection threshold at ΔRn = 0.3 (this value fits the exponential amplification phase). Dissociation curves were required to have a single peak to ensure primers specificity. The cycle thresholds (CTs) were normalized using 3 olfactory tissue-specific reference genes (i.e. *Adcy3*, *Gap43* and *Gng8*). When knockout mouse lines for one of these genes were analyzed (i.e. *Adcy3^flox/flox^;Gng8^wt/cre^* or *Gng8^del/del^*), the corresponding gene was omitted and the data was normalized using the two remaining genes. Triplicates with outliers with a standard deviation of their normalized CTs superior to 0.25 were manually discarded. Normalized CT values were converted to relative quantities (Rq) by dividing each sample’s CT to that of a randomly chosen control sample. To calculate the mRNA levels of exposed mice compared to control mice, the relative mRNA quantity of each exposed sample was divided by the mean of control samples. The statistical significance of the differences in Rq values between the groups was tested using a linear model in R, with the individual Rq values of each sample as the response variable and the group labels (i.e. exposed and non-exposed) as the predictor variable. Each gene was tested separately, and all tests were adjusted for multiple comparisons using the false discovery rate method.

##### Exon-intron split analysis

Using the STAR read aligner tool (v.2.7.0 ; (Dobin *et al*., 2013)), reads from the FACS-seq and bulk RNA-seq experiments were mapped to the *Mus musculus* Ensembl transcriptome reference (GRCm38 from Ensembl). The annotation file used for this analysis only contained protein coding gene annotations. Gene expression quantification was carried out using featureCounts (Liao *et al*., 2014) version 1.6.3. To quantify intronic reads for a specific gene, we subtracted the reads mapped on the exon from the reads mapped to the entire transcript. Transcriptional downregulation of intronic and exonic features in exposed mice was tested by a two sample independent T test using a linear model in R adjusted for FDR multiple comparisons.

#### In situ hybridization

Mice were exposed to odorants for 1 hour, after which they were euthanized. Heads were placed in 10% formalin, purged of gaz, left overnight at 4°C, transferred to 15% sucrose for 12 hours, followed by 30% sucrose for 12 hours. They were embedded in OCT and frozen. The main olfactory epithelium was cut in 16–18 μm coronal sections with a cryostat-microtome. Slides were conserved at −80 °C until use. RNA probes were designed to have a maximum identity with aspecific targets of 80% over a 100- bp window. Primers to amplify the probe for *Olfr171* were: AGTGCCTTCTCTTGGCAGT (forward) and GAGTGTGGGTGTCAGGATGG (reverse). The probe was transcribed with fluorescein-labeled UTP using the Roche RNA Labeling Kit and In-Vitro Transcription Kit following the manufacturer’s protocol. Slides were post-fixed in 10% formalin for 15 minutes, and washed for 3 minutes in PBS. Slides were incubated in 0.1% H2O2 for 30 minutes and then washed twice in PBS for 3 minutes. Slides were then treated with 10 μg/ml proteinase K in TE for 5 minutes, followed by an incubation in 10% formalin for 10 minutes and washed in PBS for 3 minutes. 0.2 M HCl was then added to the slides for 10 minutes, followed by a 3 minutes PBS wash. Then the slides were pre-incubated in 0.1 M triethanolamine HCl, pH 8 for 1 minute and incubated in 0.1 M triethanolamine HCl with acetic anhydride for 10 minutes, followed by a 3 minutes PBS wash. Probes were denatured for 7 minutes at 70 °C and diluted 1:400 in 50% formamide, 10% dextran sulfate, 1 μg/μl tRNA and 1× Denhardt’s solution in nuclease-free water. Slides were incubated in the hybridization buffer for 14–18 hours at 65 °C. Slides were washed 2× 30 minutes at 65 °C and 1× 30 minutes at 20–25 °C in 1× SSC, 50% formamide, 0.1% Tween-20, H2O DEPC. They were then pre-incubated 30 minutes with 1× MABT with 2% Blocking Reagent (Roche, ref. 11 096 176 001). Roche Anti-Fluorescein POD (Fab fragments, ref. 11426346910) was diluted 1:200 in pre-incubation mix and slides were covered with the antibody solution for 30 minutes. Slides were washed 3× 5 minutes in TNT (150 mM NaCl, 100 mM Tris, HCl to pH 7.5 in 10 L, 0.05% Tween-20), treated with PerkinElmer Biotinylated Tyramide 1:50 in Amplification Diluent for 30 minutes, and washed 3× 5 minutes in TNT. Finally they were treated with Alexa-488–labeled Streptavidin (Life Technologies) 1:100 in PerkinElmer Amplification Diluent for 30 minutes, washed 3× 5 minutes in TNT and then incubated with PBS. Fluorescence intensity was assessed by measuring the total fluorescence of a disk with an area of 1.8 μm^2^ in diameter that comprised the transcription foci using the ImageJ software. Fluorescence intensity ratios were calculated by dividing the fluorescence level of each exposed neuron to the mean fluorescence of control neurons that had been processed in parallel.

## Supplementary figures and tables

**Figure S1.**
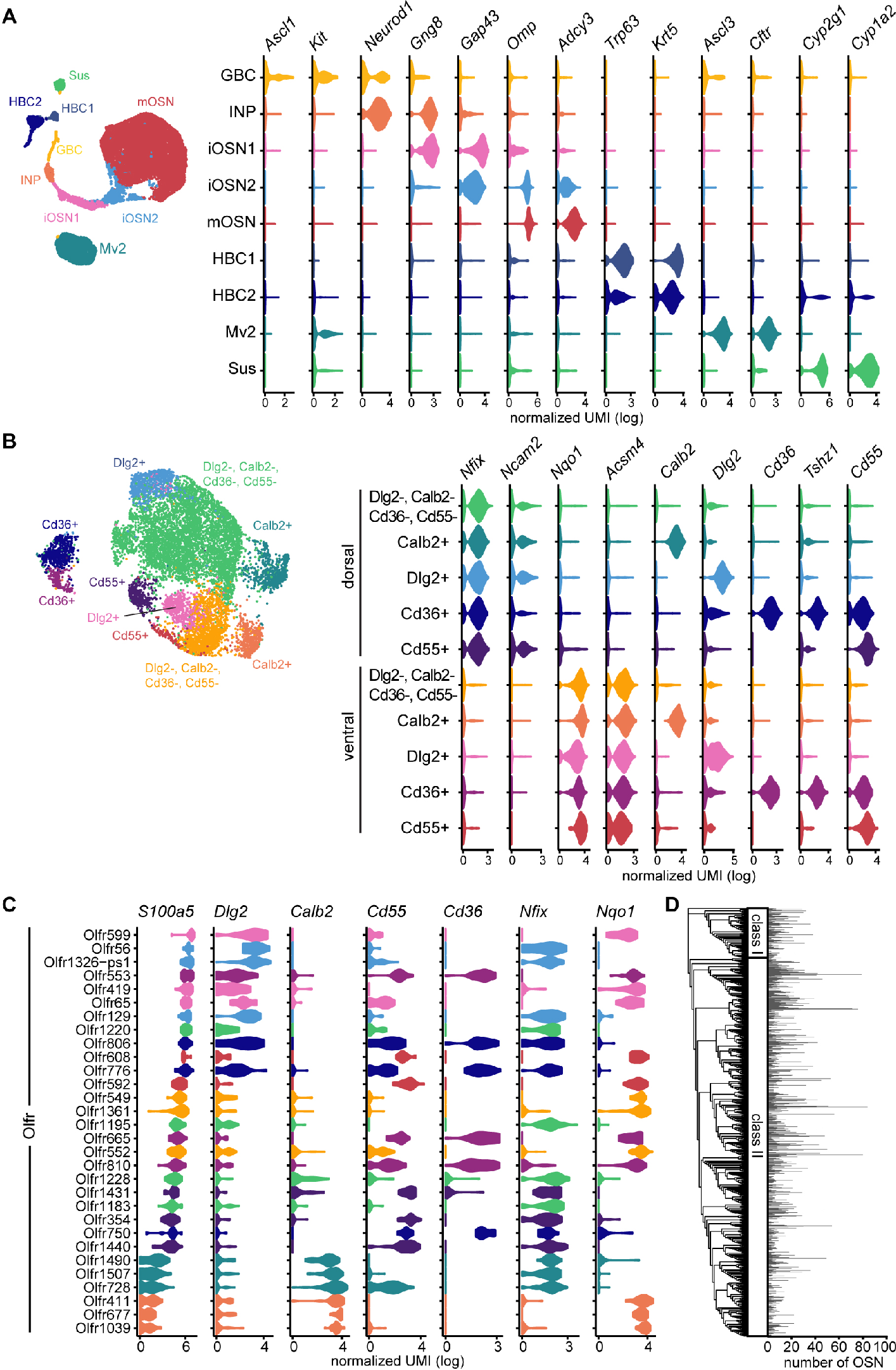
Cluster-specific markers. (A) Violin plots showing cluster-specific distribution of selective marker genes characteristic of the different cell types populating the nasal cavity (log normalized UMI). (B) Violin plots showing cluster-specific distribution of selective marker genes characteristic of the identified mature olfactory sensory neuron (mOSN) subclusters (log normalized UMI). (C) Violin plots showing cluster-specific distribution of selective marker genes characteristic of the identified mOSN subclusters (log normalized UMI) in various olfactory neuron sensory (OSN) populations. OSN populations are ordered by their mean expression of *S100a5*. The color of each violin plot indicates the cluster to which the majority of the cells from the given population pertain. D) Bar plot showing the amount of cells per OSN population composed of at least 3 cells. Olfactory receptor genes are phylogenetically organized.

**Figure S2.**
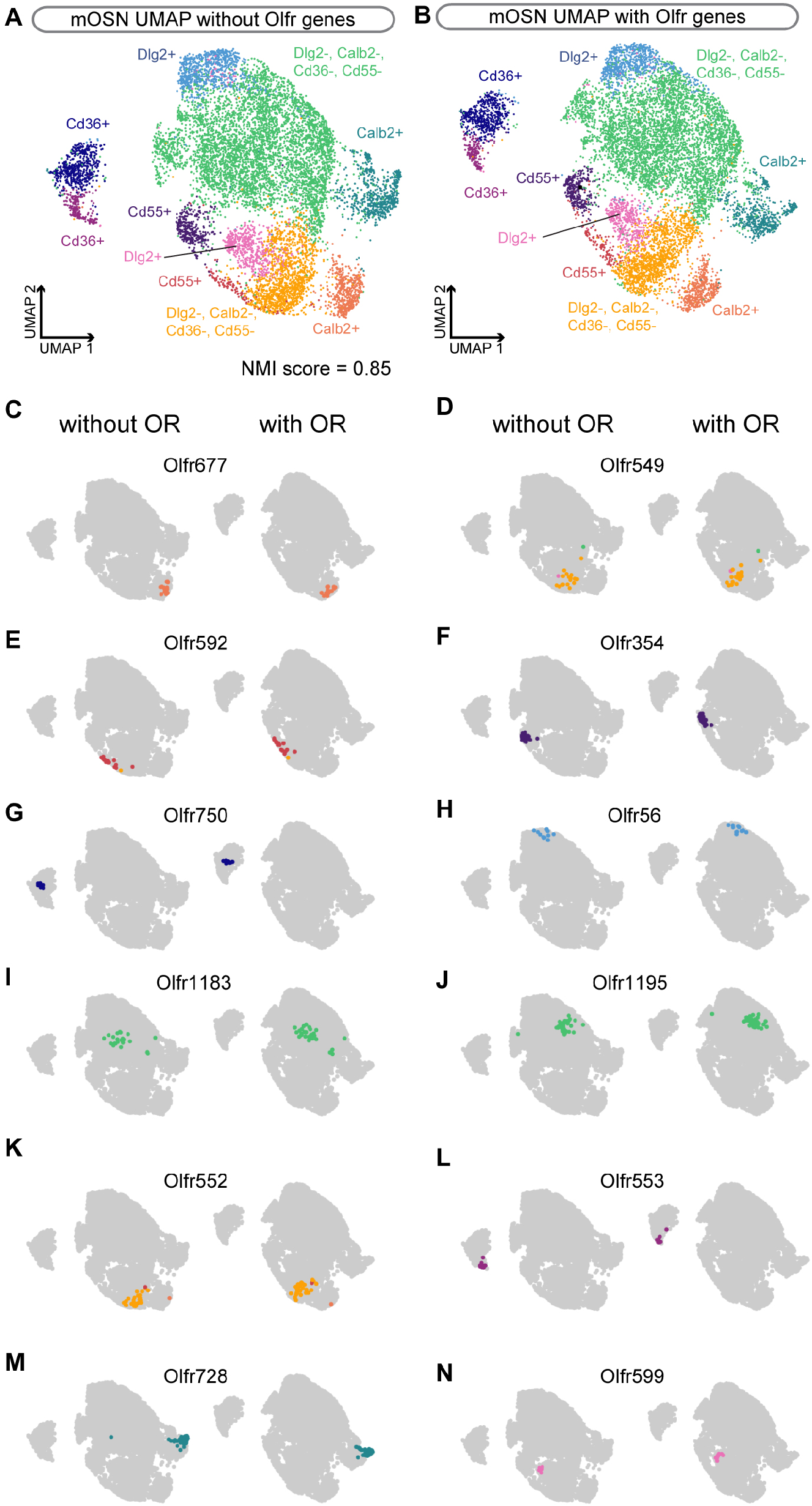
Clustering of OSN populations without odorant receptor data. (A-B) Visualization of MOE cell clusters on a *UMAP plot* computed (A) with or (B) without the inclusion of olfactory receptor genes in the count dataset. The normalized mutual information (NMI) score indicates the clustering similarity between (A) and (B). (C-N) Visualization of the dispersion of OSN populations on the UMAP plot reported in (A) (left) and (B) (right).

**Figure S3.**
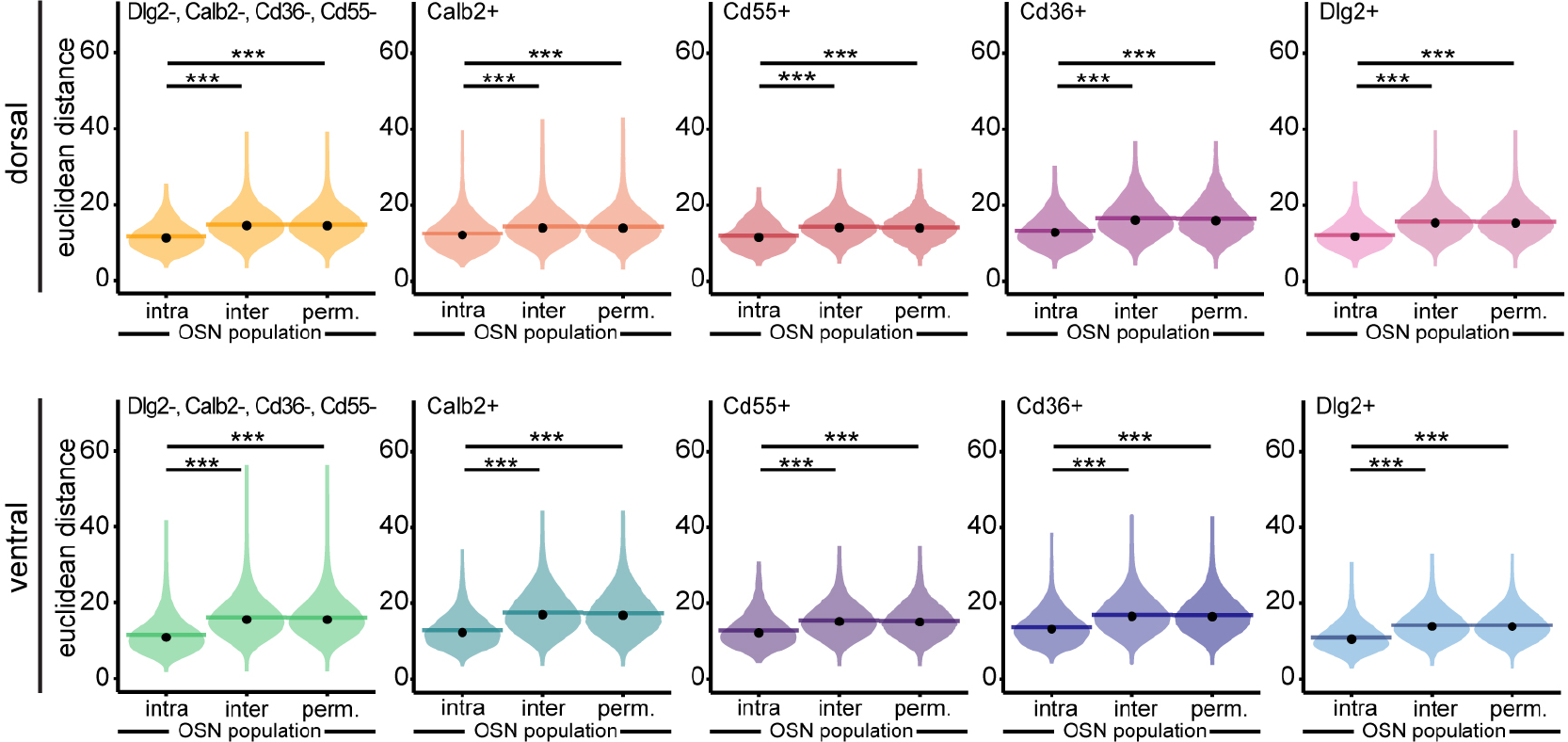
Transcriptomic Euclidian distances between OSNs. (A-C) Violin plots showing, per cluster, the density distribution of transcriptomic Euclidean distances (computed on the first 19 PCs) between pairs of OSNs expressing the same receptor (intra) or different receptors (inter). *perm.* corresponds to the distribution of Euclidean distances between pairs of OSNs expressing the same receptor after permutation (n = 1000 permutations) of receptor identities prior to distance calculation (see methods). Horizontal bars correspond to mean values and dots correspond to median values. \**p*<0.05, \*\**p*<0.01, \*\*\**p*<0.001, Wilcoxon rank test.

**Figure S4.**
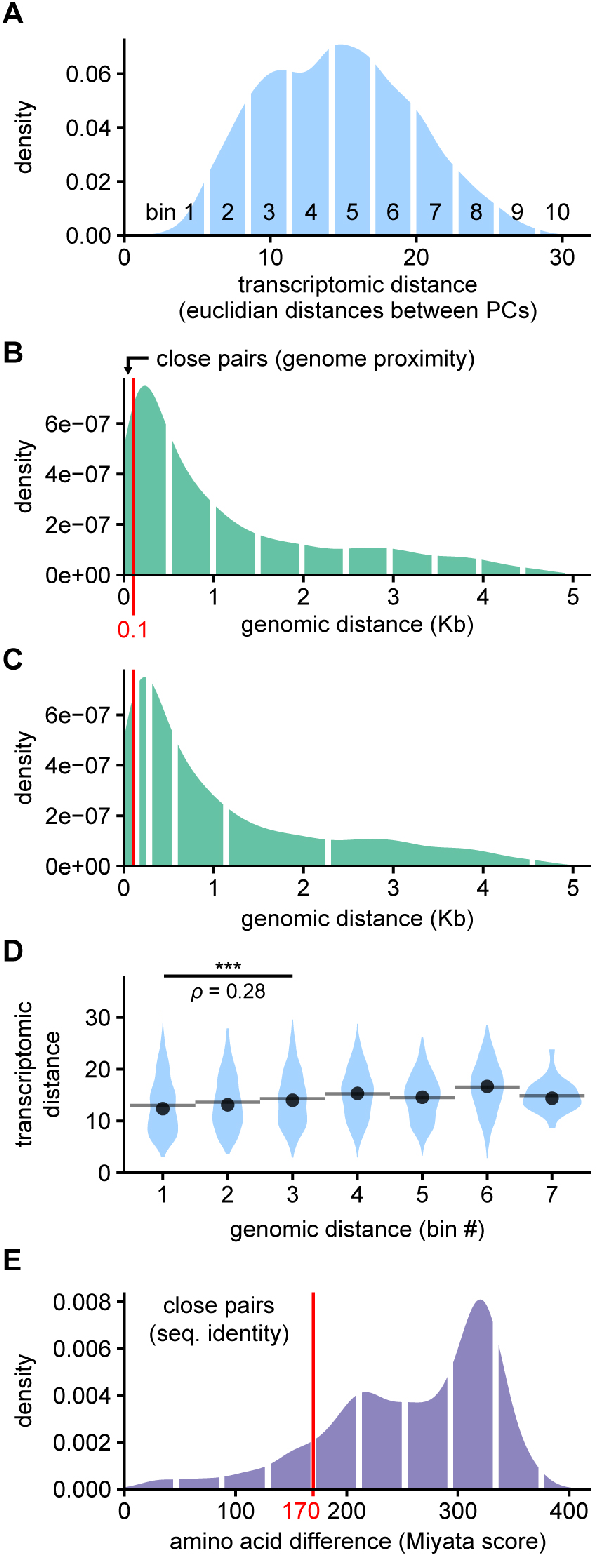
Density distributions of pairwise distance metrics. (A) Density distribution of pairwise transcriptomic distances for all pairs of OSN population expressing ORs from the same class and the same Olfr gene cluster. White lines delineate bins which cut the value range in 10 windows of equivalent width. (B) Density distribution of pairwise genomic distances for all pairs of ORs from the same class and the same Olfr gene cluster. White lines delineate bins, which cut the value range in 10 windows of equivalent width. The red line indicates the 95^th^ percentile of all intergenic distances between adjascent Olfr genes, which is the maximum genomic distance of a pair to be considered close in terms of genomic proximity. (C) Same distribution as in (B) but with a different bin definition. In this case, the right limit of the first bin is defined as four times the average intergenic distance between adjascent Olfr genes. The subsequent limits are calculated as multiplication by 2 of the precedent value. (D) Pairwise transcriptomic distance distribution for each range of pairwise genomic distance values defined by the binning described in D. Spearman’s rank correlation (ρ) was calculated for values comprised in the 3 first bins: ρ=0.28, \*\*\**p*<0.001, n=1792 pairs. (E) Density distribution of pairwise amino acid differences for all pairs of ORs from the same class and the same Olfr gene cluster. White lines delineate bins, which cut the value range in 10 windows of equivalent width. The red line indicates the 5^th^ percentile of the data, which is the threshold to separated close and distant pairs in terms of sequence identity.

**Figure S5.**
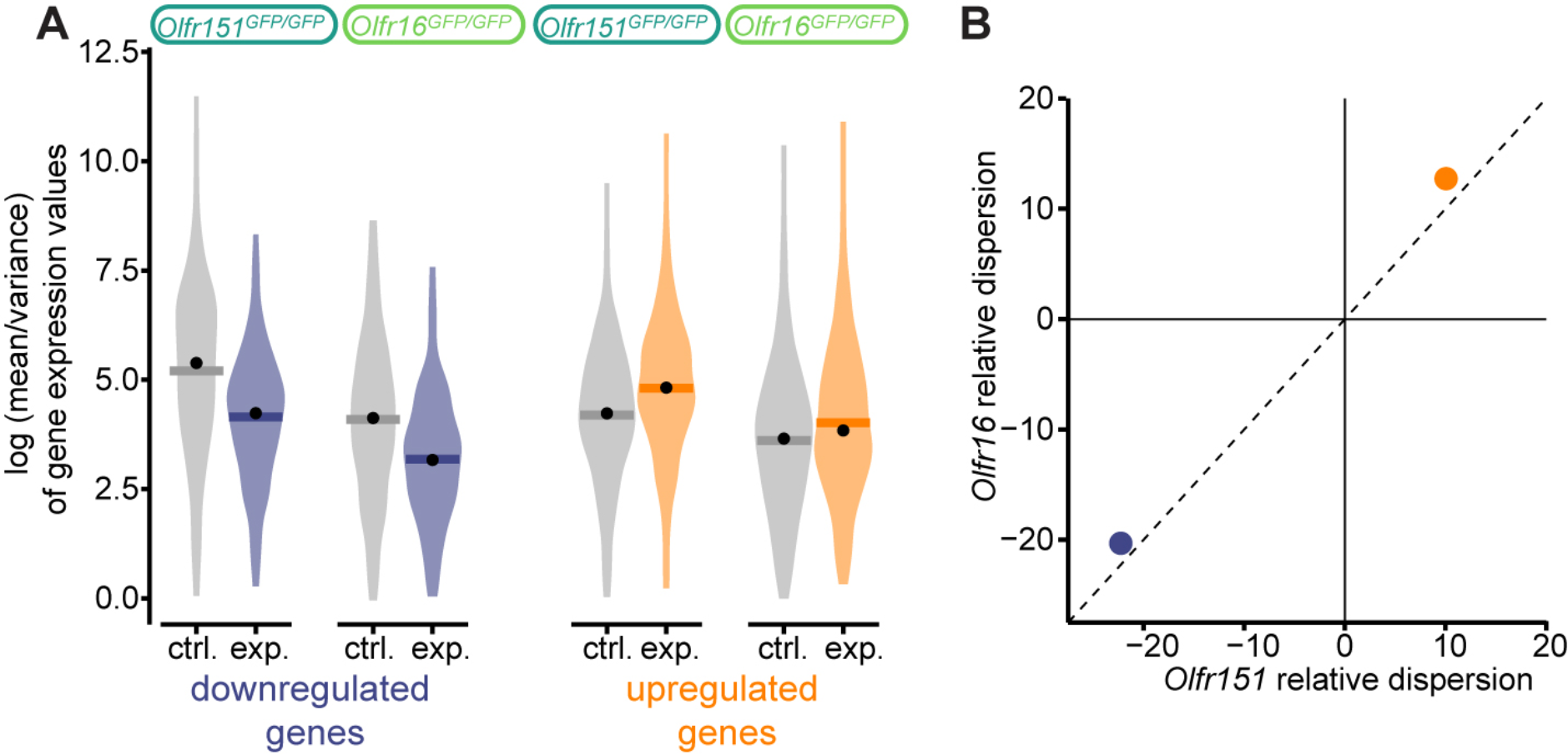
Dispersion analysis of the modulated genes in *Olfr151*(GFP/GFP) and *Olfr16*(GFP/GFP) populations. (A) Dispersion analysis in *Olfr151*(GFP/GFP) and *Olfr16*(GFP/GFP) populations. The left violin plots (purple) correspond to genes significantly downregulated after odorant exposure. The right violin plots (orange) correspond to genes significantly upregulated after odorant exposure. The y-axis represents the dispersion, calculated as log(variance/mean) per modulated gene. (B) Dispersion modulation in the exposed samples relative to the non-exposed samples, calculated as the mean of the dispersion in the exposed samples divided by the mean of the dispersion of the non-exposed samples in the modulated genes. Blue dot: relative dispersion of the downregulated genes. Orange dot: relative dispersion of the upregulated genes.

**Figure S6.**
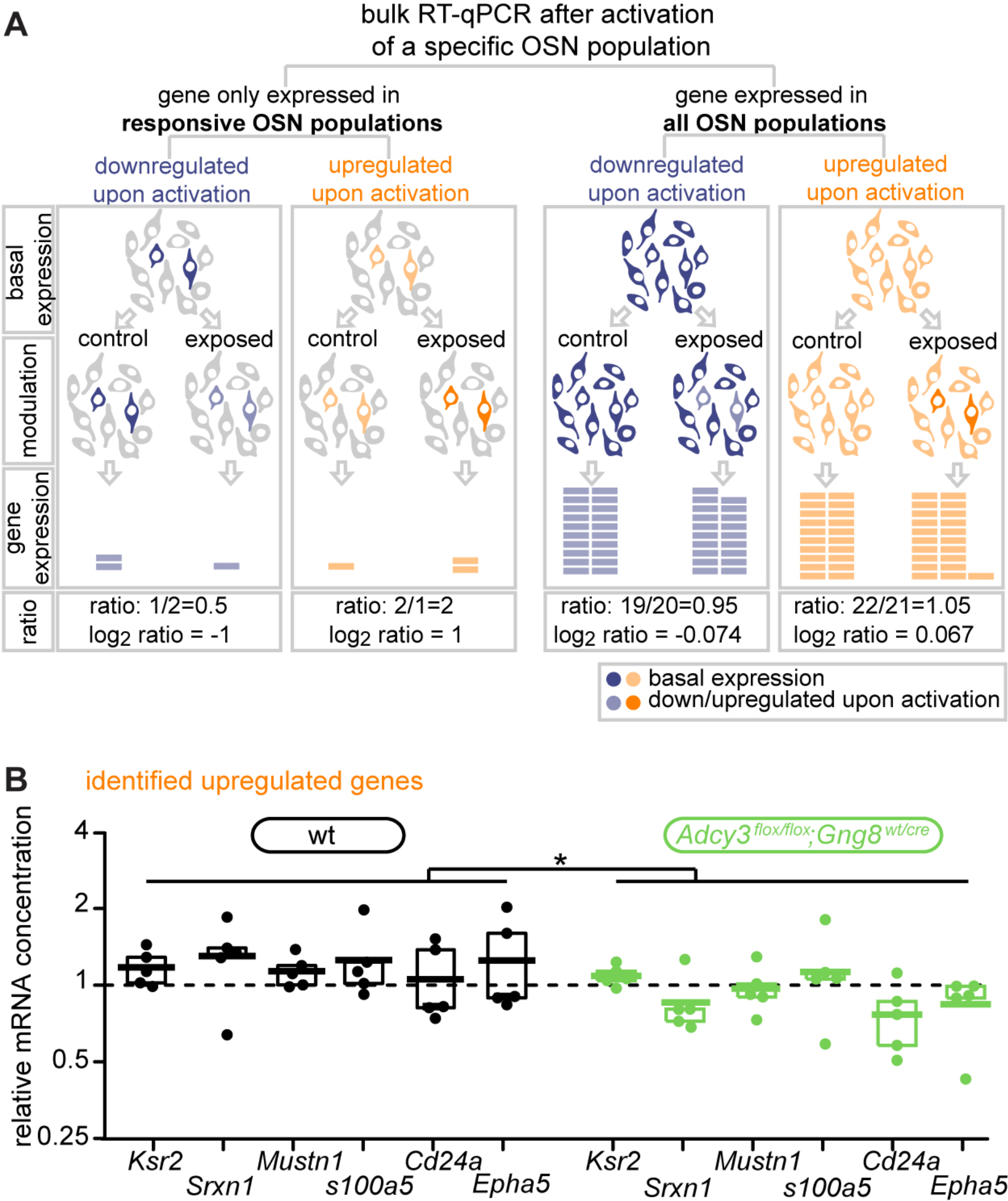
Activity-dependent upregulation of genes depends on adenylyl cyclase 3. (A) Schematic showing the significant differences of sensitivity, when using mRNA quantification of whole olfactory epithelium extracts, to identify transcriptomic modulations of genes exclusively expressed in responsive neurons or also expressed in non-responsive neurons. (B) Genes found to be upregulated in Olfr151 and Olfr16-expressing neurons after acetophenone and lyral exposure, respectively, were also found upregulated following ethyl isobutyrate exposure. This upregulation was abolished in Adcy3 null mice. \**p*<0.5, two-way RM-ANOVA.

